# Poly (ADP-ribose) induces α-synuclein aggregation in neuronal-like cells and interacts with phosphorylated α-synuclein in post mortem PD samples

**DOI:** 10.1101/2020.04.08.032250

**Authors:** Laura N. Puentes, Zsofia Lengyel-Zhand, Ji Youn Lee, Chia-Ju Hsieh, Mark E. Schneider, Kimberly J. Edwards, Kelvin C. Luk, Virginia M.-Y. Lee, John Q. Trojanowski, Robert H. Mach

## Abstract

**Background:** Poly (ADP-ribose) (PAR) is a negatively charged polymer that is biosynthesized by Poly (ADP-ribose) Polymerase-1 (PARP-1) and regulates various cellular processes. Alpha-synuclein (αSyn) is an intrinsically disordered protein (IDP) that has been directly implicated with driving the onset and progression of Parkinson’s disease (PD). The mechanisms by which αSyn elicits its neurotoxic effects remain unclear. Recent findings indicate that one of the key processes driving PD pathology are oligomeric species of αSyn. Furthermore, it is well established that the main components of Lewy bodies (LBs) and Lewy neurites (LNs) in PD patients are aggregated hyperphosphorylated (S129) forms of αSyn (pαSyn).

**Methods:** We used biochemical and immunofluorescence-based assays to explore if PARP-1 enzymatic product (PAR) drives the conversion of monomeric αSyn into aggregated assemblies. We performed quantitative measurements using in situ proximity ligation assays (PLA) on a transgenic murine model of α-synucleinopathy (M83-SNCA*A53T) and post-mortem PD/PDD patient samples to characterize PAR-pαSyn interactions. Additionally, we used bioinformatic approaches and site-directed mutagenesis to identify PAR-binding regions on fibrillar αSyn.

**Results:** Our studies show that elevated intracellular levels of PAR promote the transition of αSyn into higher molecular weight forms. We report that PAR-pαSyn interactions are predominant in pathological states. Moreover, we confirm that the interactions between PAR and αSyn involve electrostatic forces between negatively charged PAR and lysine residues on the N-terminal region of αSyn.

**Conclusions:** PAR plays a critical role in the early stages of monomeric αSyn aggregation, thereby attributing to PD pathogenesis. Based on our results, we report that PAR seeds monomeric αSyn aggregation and directly interacts with phosphorylated αSyn in conditions that are pathologically relevant to PD.

## Introduction

A characteristic feature in the pathogenesis of PD involves the accumulation of αSyn protein within the cytoplasm of brain cells^1,2^ — an event that underlies the molecular basis of PD pathology^3,4^. While the exact mechanisms associated with PD progression are unknown, it is well understood that the intracellular aggregation of αSyn is directly linked to the neurodegeneration found in PD^1^. αSyn is a protein that primarily exists as a natively unfolded soluble monomer^5^. In neurons, αSyn is believed to function in a variety of synaptic processes, including vesicle trafficking and recycling^6–8^. Depending on the environment, αSyn can undergo a variety of dynamic conformational changes, which include the formation of α-helix-rich tetramers^9^, partially folded α-helical forms (due to interactions with biological membranes), transitioning into oligomeric species, and producing toxic fibrils that are insoluble and resistant to protease activity^10^. The resulting effect of the latter is a loss in the original protein function and damage in the affected neurons^11^. In PD, αSyn accumulates into higher-order aggregates known as Lewy bodies (LBs) and Lewy neurites (LNs)^12^. The processes by which native αSyn transitions from a monomeric state to a pathogenic aggregate form are unknown. As such, identifying the underlying factors that drive abnormal αSyn assembly are vital to understand the pathogenesis of PD.

Aberrant protein aggregation has been linked to mitochondrial dysfunction and excessive production of reactive oxygen and nitrogen species (ROS/NS)^13,14^. In the last decade, extensive research has been done exploring the role of nuclear protein PARP-1 in promoting neurodegeneration^15,16^. Studies have shown that PARP-1 hyperactivation depletes NAD+, induces an accumulation of PAR, and triggers mitochondrial damage in AD^17^, HD^18^, ALS^19^, ischemic brains^20^, and PD^16^. PAR is primarily synthesized by PARP-1 from NAD+ in the nucleus of cells^21^; it regulates cellular processes such as modulating protein localization through covalent (aspartic, glutamic or lysine residues) and noncovalent interactions via PAR-binding motifs (PBMs) on target proteins^22^. Several lines of evidence show that increased levels of intracellular PAR promote liquid demixing and irreversible aggregation of IDPs^23^. Moreover, PAR and PARylated proteins have been shown to interact directly with pathogenic protein states, such as, Aβ^24^, TDP43^25^, and hnRNP-A1^26^. Thereby, affecting the aggregation kinetics of these proteins, potentiating toxicity, and promoting cell-to-cell transmission. As such, it has been suggested that the association of PAR and protein aggregates may serve as a feed-forward mechanism that amplifies neurotoxicity and drives neurodegeneration^27^.

A seminal study by Kam and colleagues^28^ revealed that αSyn preformed fibrils (PFF) increase intracellular oxidant levels which result in DNA damage and activation of PARP-1 – hyperactivation of PARP-1 leads to the intraneuronal accumulation of PAR and cell death via Parthanatos^28^. It was also reported that PAR binds αSyn PFF resulting in a more stable PFF that displays faster fibrillization kinetics and higher neurotoxicity. Although this paper presented strong experimental evidence showing an interaction between PAR and α-synuclein, several unanswered questions remained. Therefore, we felt compelled to fill this scientific gap by asking if we could establish a link between the prevalence of α-synuclein-PAR interactions and neuropathology.

To answer these research questions, we employed the use of a human neuroblastoma line overexpressing wild type αSyn wildtype (SH-SY5Y-αSyn) to gather physiologically-relevant information on the role of PAR in αSyn aggregation. We performed in situ proximity ligation assays (PLA) to gain respective insight into the pathophysiological significance of PAR-αSyn interactions. We utilized site-directed mutagenesis, immunodot blots, and molecular docking studies to elucidate the nature of these interactions. Altogether, we show that PAR promotes the pathogenic assembly of monomeric αSyn and that PAR-phosphorylated αSyn (pαSyn) interactions are predominantly observed in PD-relevant transgenic murine models of αSyn pathology and post-mortem PD/PDD patient samples.

This study provides direct insight into an early trigger of αSyn aggregation. Consistent with previous findings^28^, our results call attention to the role of PARP-1 activity (and PAR polymer) in driving PD neurodegeneration and reinforce the notion that small-molecule inhibitors of PARP-1 hold neuroprotective potential, especially in patients who are at high-risk of developing PD (i.e. patients with autosomal dominant mutations on αSyn).

## Results

### PAR polymer induces αSyn oligomerization

Kam et al.,^28^ report that treatment with *N*-methyl-*N’*-nitro-*N*-nitrosoguanidine (MNNG) – a potent pharmacological PARP activator – promotes αSyn aggregation in SH-SY5Y-αSyn and SH-SY5Y-A53T-αSyn human neuroblastoma cells. Moreover, they report that exogenous administration of PAR induces αSyn aggregation in SH-SY5Y WT and SH-SY5Y PARP-1 KO cells^28^. In our study, we set out to expand on these findings and evaluate whether PAR polymer promotes the transition of natively unstructured αSyn into a disease-associated aggregate state (Fig. 1a).

**Figure 1.**
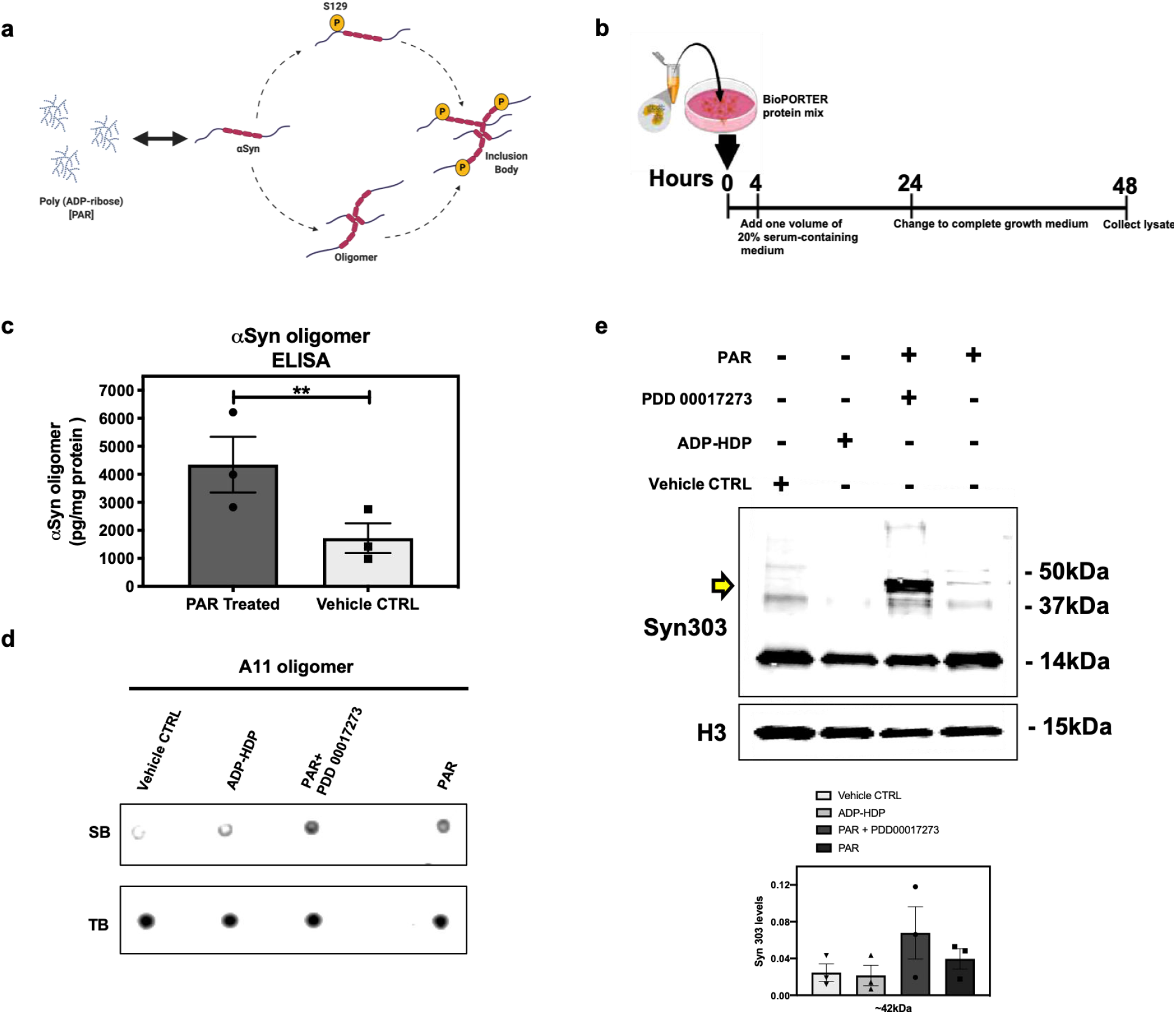
PAR seeds monomeric αSyn aggregation. (a) Proposed molecular mechanism of PAR induced αSyn aggregation (b) Experimental scheme of *BioPORTER*-mediated transfection of PAR polymer into SH-SY5Y-αSyn neuroblastoma cells. (c) Human αSyn oligomer-specific ELISA on cell lysates from SH-SY5Y-αSyn cells treated with 50 nM PAR vs *BioPORTER* alone (vehicle control). Bars represent mean ± SEM. Student’s one-tailed t test (n = 3). **P = 0.003. R^2^ = 0.53. d.f. = 10. (d) Representative Immunodot blot of an oligomer-specific antibody (A11) used to measure specific binding (SB) i.e. oligomeric content in PAR, PAR + 1 μM PDD00017273 (PARGi), and ADP-HDP treated cell lysates vs. *BioPORTER* alone (vehicle control). A total protein stain (Revert 700) was used to assess total binding (TB). Experiments were repeated independently three times (n = 3). (e) Representative Western blot (top) and semi-quantitative analysis of higher molecular weight (~ 42kDa) αSyn aggregation levels using conformation-specific Syn 303 antibody (bottom). Bars represent means ± SEM. One-way ANOVA (n = 3). P = ns. Histone 3 (H3) was used as a loading control.

Commonly used pharmacological agents that induce PARP-1 hyperactivation also trigger significant genotoxic stress^29–31^. Therefore, to characterize the role of PAR in seeding αSyn aggregation, we employed the use of a protein transduction system, *BioPORTER,* to deliver a physiologically relevant dose of PAR polymer (50 nM) into SH-SY5Y-αSyn cells (Fig. 1b). The rationale for the use of *BioPORTER –* instead of a genotoxic agent like MNNG^29^ – was to develop a neuronal-like cell model that recapitulated the effects of PARP-1 hyperactivation (i.e. elevated PAR) in a genomically stable setting. To determine if PAR itself affects cell viability, we performed cytotoxicity assays on SH-SY5Y-αSyn cells. Results from these assays showed that PAR administration did not lead to significant cell death (Extended Data Fig. 1a).

To assess if PAR increases intracellular superoxide levels, we performed live cell imaging 48 h post PAR/*BioPORTER* delivery in transiently transfected Hela cells that were overexpressing an EGFP-αSyn-A53T fusion protein (Extended Data Fig. 1b). Superoxide levels were detected with MitoSOX™ Red (a superoxide indicator reagent) and imaged via fluorescence microscopy. Results from these studies showed that PAR treatment increased intracellular superoxide levels in Hela cells when compared to *BioPORTER* alone (Extended Data Fig. 1b).

To obtain a quantitative measure of αSyn oligomerization, we used an enzyme-linked immunosorbent assay (ELISA), specific for human αSyn oligomers, to evaluate if elevated levels of PAR in SH-SY5Y-αSyn cells promote the conversion of monomeric αSyn into oligomeric forms. Results from the ELISA (Fig. 1c) indicate that cell lysates from PAR treated samples had higher αSyn oligomeric signal compared to vehicle controls.

To further validate our ELISA data, we set up an immunodot blot experiment, whereby, cell lysates were spotted onto a membrane and assayed for the presence of amyloid oligomers using an A11 antibody that has been previously validated by our group^32^ and others^33,34^ and is specific for prefibrillar oligomers ranging in size from 40 – 75 kDa^34^ (Fig. 1d, Supplementary Table 1). To investigate if the increase in A11 oligomer signal (48 h post-treatment) was PAR-specific, we used a small molecule PARG inhibitor (PDD 00017273)^35^ to increase endogenous PAR levels. PARG is an enzyme that regulates intracellular PAR levels via its exo- and endoglycosidase activities^36^. Thus, to reduce PAR catabolism, we pre-treated SH-SY5Y-αSyn cells with 1 μM PARG inhibitor PDD 00017273, 1 h prior to *BioPORTER* delivery of 50 nM PAR. Additionally – and in parallel – we used *BioPORTER* to deliver 50 nM of adenosine diphosphate (hydroxymethyl)pyrrolidinediol (ADP-HDP)^37^ into SH-SY5Y-αSyn cells to assess if the stable NH-analog of ADP-ribose was sufficient to induce intracellular αSyn aggregation (Fig. 1d). Results from the A11 immunodot blot assay showed higher oligomeric signal in the PAR and PAR + PARGi samples, compared to vehicle control and ADP-HDP treated samples (Fig. 1d); thus, corroborating our αSyn oligomer ELISA data (Fig. 1c).

Since misfolded αSyn seeds and drives pathology in cell and animal models of PD^38,39^, we performed biochemical analysis to test for the presence of higher molecular weight forms of αSyn. For these experiments, we used a conformation-selective αSyn antibody (Syn303) which recognizes total and misfolded synuclein^40,41^ (Fig. 1e, Supplementary Table 1). Results from these experiments showed an increase in band intensity for a higher molecular weight form of αSyn (~ 42 kDa) in the PAR + PARGi treated samples 48 h post treatment (Fig. 1e). Moreover, these results confirm that continuously elevated levels of PAR promote the formation of higher molecular weight αSyn aggregates.

Recent evidence suggests that αSyn is a DNA binding protein that interacts with γH2AX and PAR on sites of DNA damage^42^. Therefore, we sought to determine if elevated levels of PAR can lead to DNA damage. We immunoblotted for γH2AX and observed an increase in γH2AX band intensity in the PAR and PAR + PARGi treated samples when compared to vehicle control and ADP-HDP treated samples (Extended Data Fig.1c). This data suggests that PAR alone can trigger DNA damage. Since PARP-1 activity accounts for the majority of PAR in the cell^43^, it stands to reason that PARP-1 hyperactivity (i.e. elevated PAR formation) promotes further DNA damage. Thus, a plausible scenario arises, whereby PARP-1 hyperactivation^44,45^ promotes the interactions of PAR and αSyn at sites of DNA damage.

In summary, our results suggest that elevated PAR levels, while not directly toxic, can lead to oxidative stress and DNA damage. Furthermore, we show that PAR itself promotes the transition of monomeric αSyn into higher molecular weight (i.e. oligomeric) forms.

### PAR colocalizes with phosphorylated (S129) αSyn Aggregates

In physiological settings, approximately 4% of soluble αSyn is phosphorylated at amino acid residue S129 (pαSyn)^46,47^. Correlations have been established between pαSyn status and pathological conditions^48,49^. In PD patients, αSyn found within inclusions (LBs, LNs) tend to be hyperphosphoryated at S129. In LBs, it is estimated that up to 90% of αSyn is phosphorylated at S129^47^. pαSyn is observed in other synucleinopathies (neurodegenerative diseases characterized by abnormal accumulation of αSyn aggregates) including dementia with LBs (DLB)^50^ and multiple system atrophy (MSA)^51^. In addition, increased levels of pαSyn has been reported in PD-like transgenic murine models^6^.

Here, we asked whether PAR interacts directly with pαSyn aggregates. To address this question, we used our previously described cell model (Fig. 1b) which mimics PARP-1 hyperactivation and introduced either: PAR, ADP-HDP or *BioPORTER* alone (vehicle control). After a 48 h incubation, the cells were immunostained with an antibody directed towards pαSyn. Using fluorescence microscopy, we identified pαSyn inclusions (~ 1 μm length) in the cytoplasm of PAR treated cells (Fig. 2a,b). We also noted that PAR signal overlapped with ~ 60% of these cellular pαSyn inclusions when co-immunostaining with a PAR-specific antibody (Fig. 2c); antibody information can be found in Supplementary Table 1.

**Figure 2.**
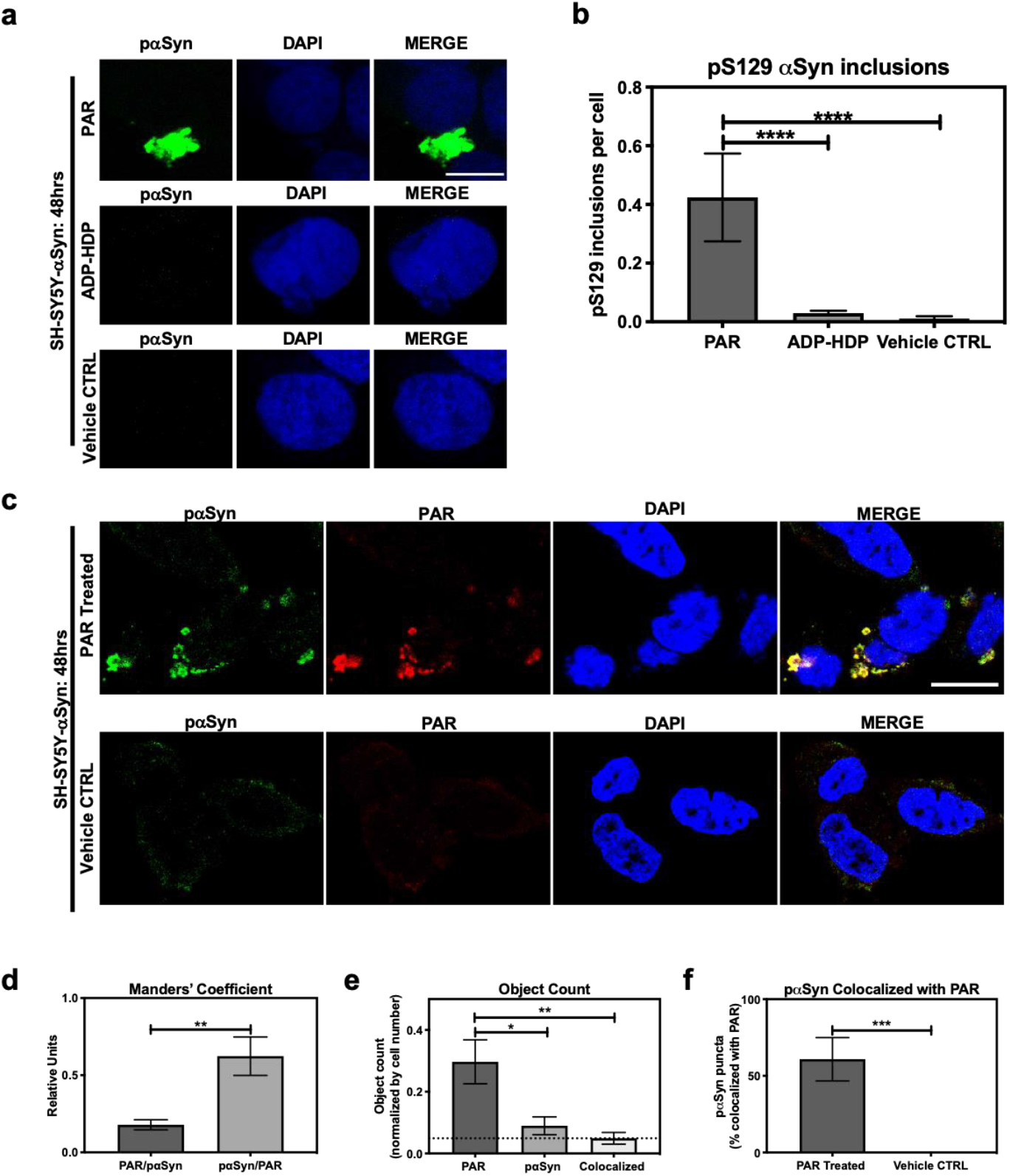
PAR promotes the formation of αSyn inclusion aggregates. (a) Representative immunostain of pαSyn (green) and DAPI (blue) in SH-SY5Y-αSyn cells 48 h post treatment with either 50 nM PAR or ADP-HDP vs. *BioPORTER* alone (vehicle control). (b) Quantification of pαSyn inclusions (aggregates larger than 1 μm) in PAR treated, ADP-HDP treated, and *BioPORTER* alone (vehicle control) samples. Bars represent means ± SEM. Two-way ANOVA followed by Tukey’s post hoc test (n = 3). ****P<0.0001. (c) Representative IF immunostain of pαSyn (green), PAR polymer (red) and DAPI (blue) in SH-SY5Y-αSyn cells 48 h post treatment with either 50nM PAR vs. *BioPORTER* alone (vehicle control). Scale bar 10 μm. (d) Manders’ overlap coefficient analysis (where an overlap coefficient of 0.5 implies that 50% of both objects, i.e. pixels, overlap) between PAR over pαSyn inclusions (PAR/pαSyn) against pαSyn inclusions over PAR (pαSyn/PAR). Bars represent means ± SEM. Student’s two-tailed t test (n = 3). **P < 0.002 (e) Object count analysis normalized by DAPI count i.e. cell number. Bars represent means ± SEM indicating total object counts for PAR, pαSyn inclusions, and total colocalized objects. The graph indicates that most pαSyn immunostain was colocalized with PAR, but a significant amount of PAR was not colocalized with pαSyn. Two-way ANOVA followed by Tukey’s post hoc test (n = 3). *P < 0.03, **P < 0.002 (f) Colocalization analysis comparing the number of pαSyn inclusions colocalized with PAR immunostain in PAR treated and *BioPORTER* alone (vehicle control) samples. Images were captured using Zeiss LSM 710 confocal (40x/1.4 Oil) microscope. Bars represent means ± SEM. Student’s two-tailed t test (n = 3). ***P < 0.0005. Scale bar 10 μm.

Our studies show that PAR-mediated αSyn aggregation results in the appearance of cytosolic pαSyn inclusions (Fig. 2a,c) by 48 h. By contrast, these pαSyn inclusions were not observed in the ADP-HDP treated or vehicle control samples (Fig. 2a,b). In addition, quantification of co-immunostained samples using image processing software, indicate that while over half of the pαSyn inclusions were colocalized with PAR signal (Fig. 2d,f), the majority of PAR signal was not colocalized with pαSyn (Fig. 2d). The latter was not surprising given the diverse roles that PAR plays in the cell^52,53^.

### PAR directly interacts with phosphorylated (S129) αSyn

To directly measure the interactions between PAR and pαSyn in our cell model, we used antibodies against PAR^54^ and pαSyn^55^ (Supplementary Table 1) and employed the use of an in-situ PLA^56^. The utilization of PLA allowed us to record the prevalence of PAR-pαSyn interactions with greater sensitivity and specificity when compared to traditional immunoassays.

A time course experiment on our cell model, showed a clear time-dependent difference in PLA signal by 48 h between the PAR and PAR + PARGi treated samples in comparison to the ADP-HDP treated and vehicle control samples (Fig. 3a,b); earlier time points (i.e. 4 h and 24 h) also showed higher PLA signal for the PAR and PAR + PARGi treated samples (Extended Data Fig. 1d,e). Our data showed an initial increase in PLA signal for ADP-HDP treated samples at 4 h (Extended Data Fig. 1d), however, after 24 h the signal fell below detection for this treatment condition (Extended Data Fig. 1e). A potential reason for the drop in PLA signal after 24 h – for the ADP-HDP treated samples – could be attributed to the degradation of ADP-HDP by phosphodiesterases in the cell^57^.

**Figure 3.**
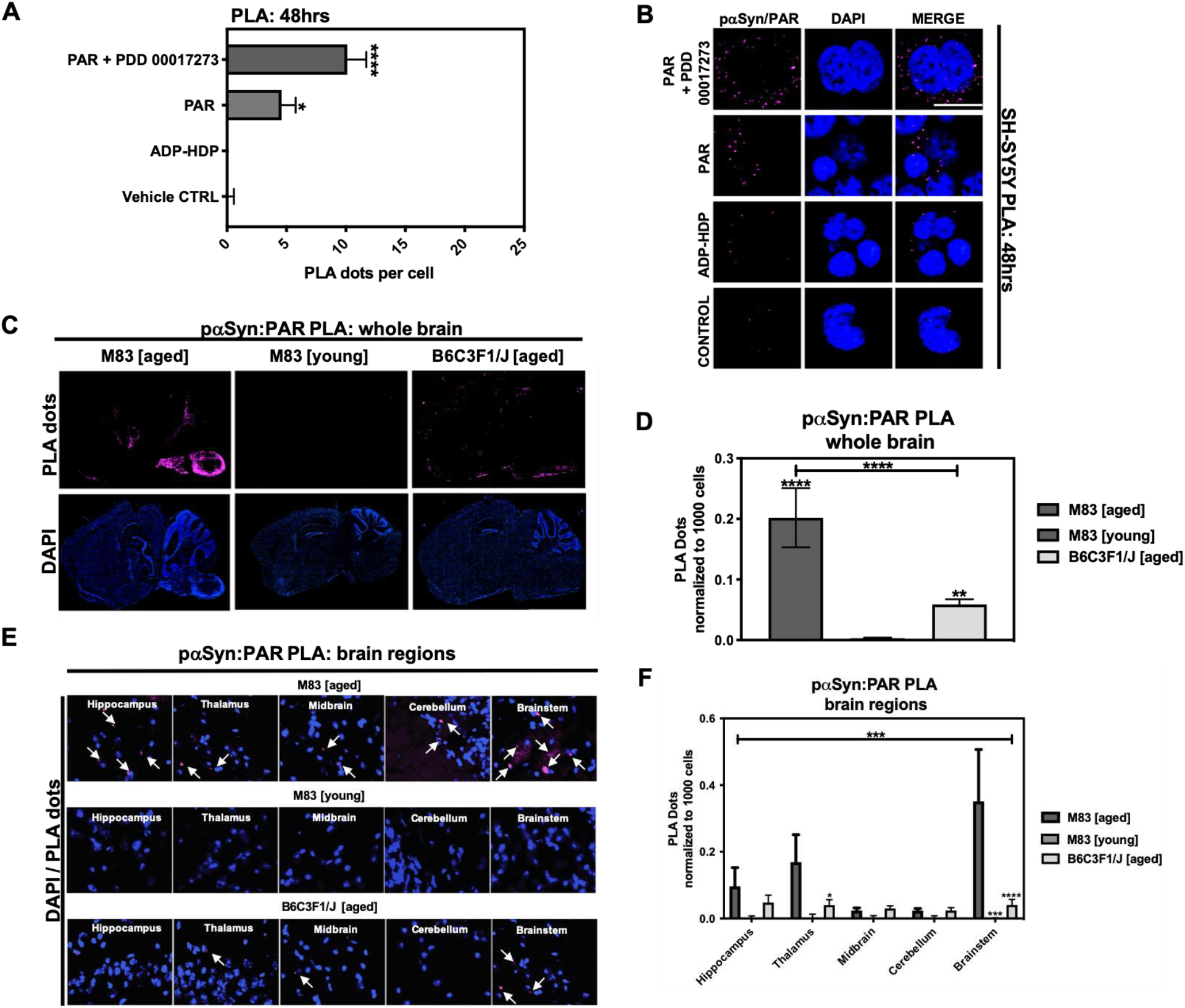
PAR interacts with phosphorylated (S129) αSyn in pathological settings. (a) Quantification from proximity ligation assays (PLA) measuring pαSyn and PAR interactions in SH-SY5Y-αSyn cells 48 h post treatment with either PAR, PAR + 1 μM PDD00017273 (PARGi) or ADP-HDP vs. *BioPORTER* alone (vehicle control). Bars represent means ± SEM. Two-way ANOVA followed by Tukey’s post hoc test (n =3). *P < 0.03, ****P < 0.0001 (b) Representative DAPI and PLA ROI images showing PLA dots (pink); these dots indicate direct interactions between pαSyn and PAR in PAR treated, ADP-HDP treated and *BioPORTER* alone (vehicle control) samples. Scale bar 10 μm. (c) Representative DAPI (bottom panel) and PLA (top panel) whole brain images from M83 Tg aged, M83 Tg young, and B6C3F1/J aged mice (n = 3 mice per group). (d) Quantification of whole brain PLA levels in M83 Tg aged, M83 Tg young and B6C3F1/J mice (n =3). Each bar represents means ± SEM. One-way ANOVA. **P < 0.002, ****P < 0.0001 (e) Representative PLA staining (white arrows) of ROIs obtained from 20x merge images. (f) Quantification of different brain regions in M83 Tg aged, M83 Tg young, and B6C3F1/J aged mice. Images were captured using Zeiss Axio Widefield (20x/0.8) microscope. Bars represent means ± SEM. Two-way ANOVA followed by Tukey’s post hoc test (n = 3). **P < 0.002, ***P < 0.0002, ****P < 0.0001.

In accordance with our previous findings, not only does PAR polymer promote the oligomerization of αSyn, it also directly interacts with pαSyn inclusions. The evidence from our cell model, suggests that PAR has a *dual* role in the aggregation pathway of αSyn, whereby it functions as a molecular seed that nucleates monomeric αSyn (resulting in higher molecular weight forms of αSyn) and perhaps stabilizes pαSyn inclusions, as evidenced by our PAR-pαSyn colocalization and PLA studies.

### PAR and pαSyn interactions are prevalent in PD-like transgenic mouse models of α-synucleinopathies

In order to assess if PAR-pαSyn interactions are present in αSyn pathology, we used a transgenic (Tg) murine model of α-synucleinopathy (M83 SCNA*A53T) that develops a PD-like phenotype with age^58^. The Tg murine line (M83) used in this study overexpresses a form of human αSyn that harbors a point mutation at amino acid residue 53 (A53T); this point mutation has been directly implicated in familial early onset PD^59^. Information on the animals used in this study can be found in Supplementary Table 2.

The Tg mice were separated into two groups: M83 Tg young (less than 12 mo) and M83 Tg aged (12 mo or older). To control for age-related effects, we also used a non-transgenic murine line (B6C3F1/J) in our studies (Supplementary Table 2). Following euthanasia, murine brains were dissected and hemisected in the sagittal plane. Immunostaining for endogenous PAR was carried out to assess PARP-1 activity. PAR intensity (particularly in the cerebral cortex) increased with age in both M83 Tg and non-Tg mice (Extended Data Fig. 2a). Similarly, we immunostained M83 Tg and non-Tg mice to assess whole brain pαSyn expression (Extended Data Fig. 2b). Based on these experiments, we observed a remarkable increase in pαSyn expression in an M83 Tg aged (17 mo) sample – with maximum signal output measured in the brainstem and cerebral cortex regions (Extended Data Fig. 2b). In addition, we show that PAR and pαSyn staining significantly increased with age in the M83 Tg brain samples, thus suggesting an association between increased PAR and pαSyn expression and disease phenotype.

To measure the prevalence of PAR-pαSyn interactions, we performed in situ PLA on frozen brain sections from all three animal groups: M83 Tg young, M83 Tg aged and non-Tg (B6C3F1/J). To limit non-specific PLA signal, we included technical controls, whereby adjacent brain tissue sections were incubated without primary antibodies (data not shown), subsequent imaging parameters (i.e. exposure time, depth of field) were then adjusted in order to acquire detectable signal above background. To ensure consistency between experimental models, we used the same primary antibodies (anti-PAR and anti-pαSyn, Supplementary Table 1) for our cell and animal brain tissue PLA.

Results from our studies revealed that PLA signal for PAR-pαSyn was strongest in M83 Tg vs. non-Tg mice (Fig. 3c,d). Analogously, M83 Tg aged vs. M83 Tg young mice differed significantly in PLA signal (Fig. 3c,d) – with the strongest signal differential detected in the brainstem region of the M83 Tg aged group (Fig. 3e,f). Similarly, increased PLA signal was also observed in an M83 Tg aged mouse (18 mo) when using primary antibodies against PAR and total αSyn (anti-PAN-αSyn) (Extended Data Fig. 2c), thus confirming that PAR interacts with both phosphorylated and non-phosphorylated αSyn. In addition, we noted that M83 Tg brain tissue samples with the highest pαSyn expression also had the greatest PLA signal output, suggesting that pαSyn-PAR PLA signal is directly tied to the amount of pathology (i.e. pαSyn expression) present in a given sample.

Overall, our studies revealed that PLA signal was highest in anatomical brain regions most commonly associated with αSyn pathology in the M83 Tg aged group (*38*) (Fig. 3e,f); these findings are in accordance with our observations from the SH-SY5Y-αSyn cell model, which show that PAR-pαSyn interactions are prevalent in pathogenic states involving αSyn aggregation and PARP hyperactivity i.e. elevated PAR levels.

### PAR-pαSyn interactions are observed in post mortem brain tissue from PD/PDD patients

To determine the clinical relevance of PAR-pαSyn interactions in PD, we performed immunoassays and PLA on human post mortem striatum and midfrontal gyrus brain regions derived from PD and PDD (Parkinson’s Disease Dementia) patients, as well as, non-PD controls. Patient information can be found in Supplementary Table 3.

Results from these studies, revealed heterogeneity in pαSyn (Extended Data Fig. 3a) and PAR (Extended Data Fig. 3b) immunostaining for all patient samples. Overall, PD/PDD patient samples exhibited increased pαSyn expression (Fig. 4a,b), while PAR expression in the PD/PDD patient samples was more heterogeneous (Extended Data Fig. 3b). Interestingly, we observed higher signal overlap (i.e. colocalization) between PAR and pαSyn staining in the PD/PDD patient samples, as determined by Manders’ overlap coefficient. (Fig. 4d). Cumulatively, the PD/PDD patient group had higher pαSyn (Fig. 4b), PAR (Fig. 4c), and PLA (Fig. 5 a–c) signal, when compared to the non-PD controls. Moreover, PLA signal output for the PD/PDD patient group was in accordance with our PAR-pαSyn colocalization analysis.

**Figure 4.**
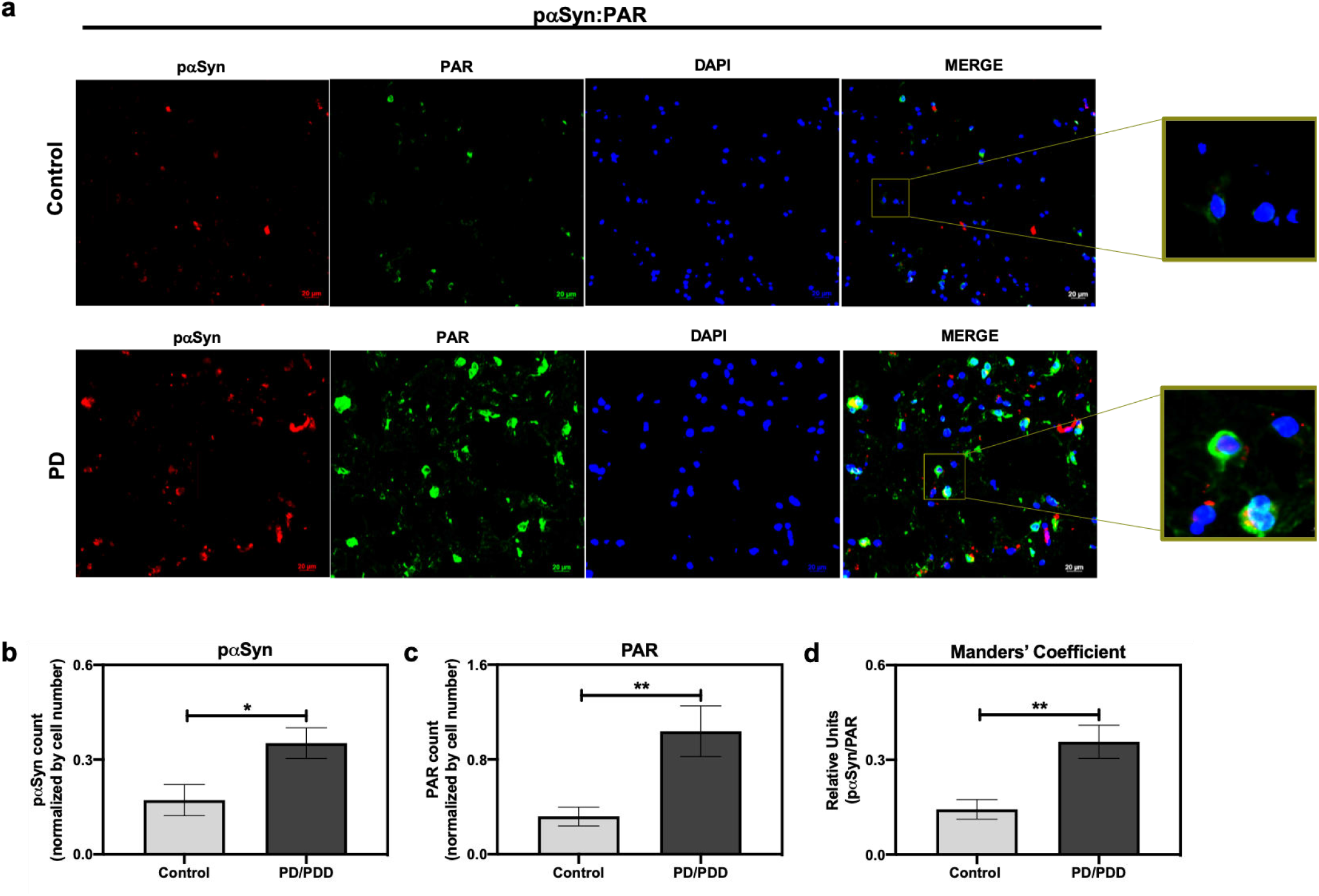
Increased S129 pαSyn and PAR levels in PD/PDD patient samples. (a) Representative IF immunostain of pαSyn (red), PAR (green), and DAPI (blue) in non-PD control (top panel, tissue ID 121111) and PD/PDD (bottom panel, tissue ID 116441) patient samples. Merge channel regions of interest (ROI) show colocalization between pαSyn and PAR staining. Scale bar 20 μm. Images were captured using Zeiss Axio Widefield (20x/0.8) microscope. (b) Quantification of pαSyn levels, normalized by DAPI count, in control vs. PD/PDD patients. (c) Quantification of PAR levels, normalized by DAPI count, in control vs. PD/PDD patients. (d) Quantification of pαSyn/PAR overlap in control vs. PD/PDD patients using Manders’ overlap coefficient. Bars represent means ± SEM. Student’s two-tailed t test (n = 3 to 4 patient samples per group). *P < 0.03, **P < 0.002.

**Figure 5.**
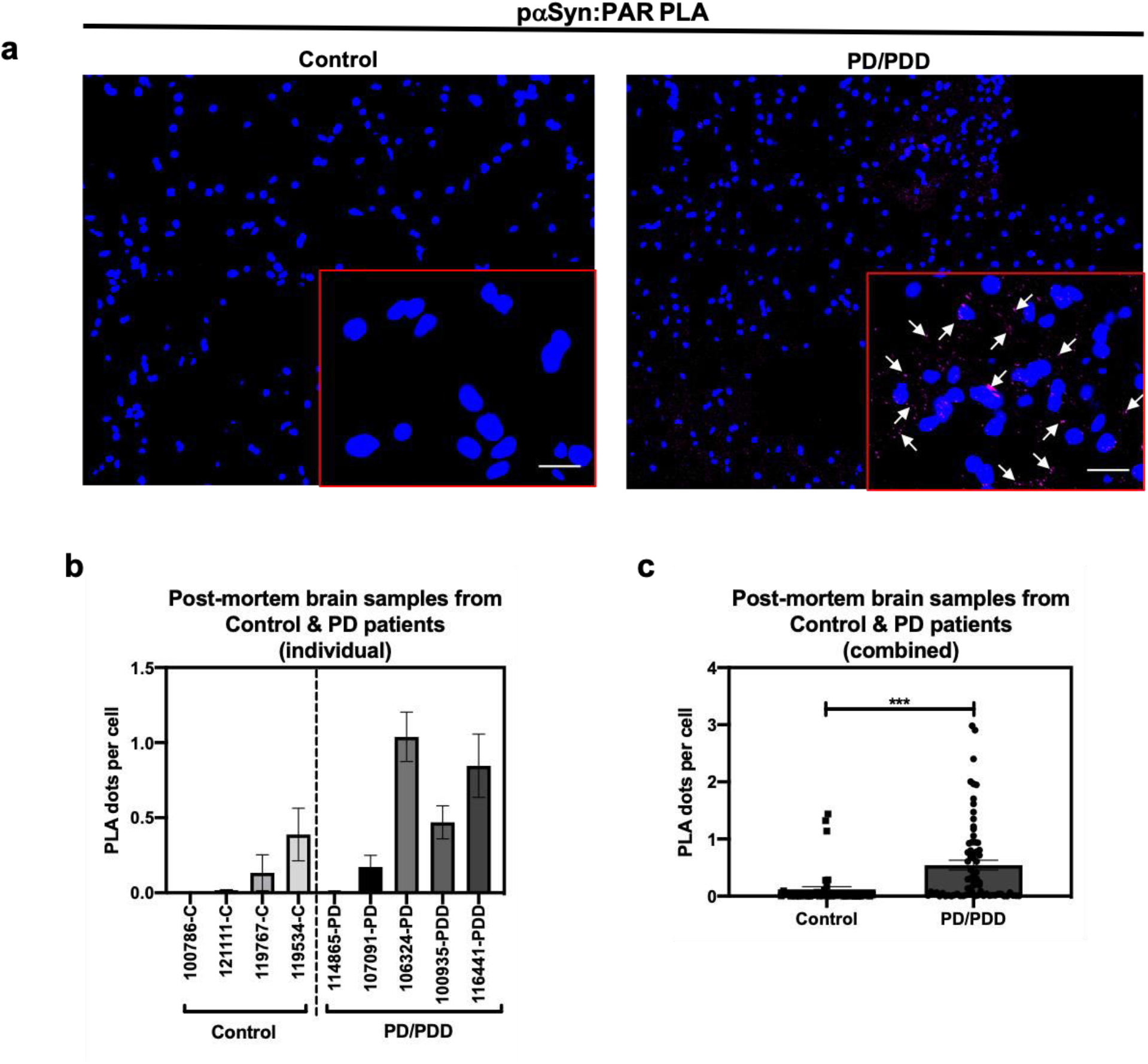
PAR predominantly interacts with pαSyn in PD/PDD post mortem patient samples. (a) Representative PLA and DAPI images in non-PD control (top panel, tissue ID 121111) and PD/PDD patient samples (bottom panel, tissue ID 116441). ROI overlaid on merge channel. ROI scale bar 10 μm. Images were captured using Zeiss Axio Widefield (20x/0.8) microscope. Representative white arrows showing PLA positive signal (b) Quantification of individual PLA dot count in non-PD control vs. PD/PDD patient samples. Bars represent means ± SEM. (c) Combined PLA analysis for all control and PD/PDD tissue samples, whereby each graphical symbol (black circle) represents the average number of PLA dots normalized to cell count (DAPI) for each field-of-view. 15 fields were captured for each patient sample. Student’s two-tailed t test (n = 3 to 4 patient samples per group). ***P < 0.0005.

To further validate our PD/PDD PLA results, we performed pαSyn-PAR PLA on cerebellum tissue samples from patients diagnosed with multiple system atrophy (MSA) – another main type of α-synucleinopathy – along with healthy cerebellum tissue controls (Extended Data Fig. 3c). We observed PLA staining patterns that closely matched pαSyn pathology in MSA (Extended Data Fig. 3 c,d). As a result, we were able to validate PLA signal in two different α-synucleinopathies.

In summary, even though we observed heterogeneous expression for pαSyn and PAR staining in the patient tissue samples, based on our analysis, we can deduce that PLA signal is higher in the PD/PDD patient group. The data obtained from these experiments, to our knowledge, is the first direct evidence showing pathologically relevant PAR-pαSyn associations on human post mortem brain tissue samples from PD/PDD and MSA patient groups. Further studies are merited in order to better understand this association as it would be of medical significance to evaluate if PAR-bound pαSyn has potential as an early PD/PDD biomarker – such a finding could have wide ranging implications for the development of disease modifying therapies for patients harboring familial PD genetic variants (i.e. A30P, E46K, and A53T).

### PAR binds αSyn via electrostatic interactions involving lysine residues

A previous study^28^ showed that PAR binds αSyn via non-covalent interactions on the N-terminal region of αSyn – thus, suggesting that the interactions between PAR and αSyn are electrostatic in nature. In order to identify the amino acid residues involved in αSyn-PAR binding, a protein alignment tool (NPS@PATTINPROT search) was used to align the native αSyn protein sequence to two published PAR binding motifs (PBM)^22,60^. We identified two sites on αSyn as potential PAR-binding sites (Fig. 6a): a site between amino acids residues 43-54 and another site between amino acid residues 48-58.

**Figure 6.**
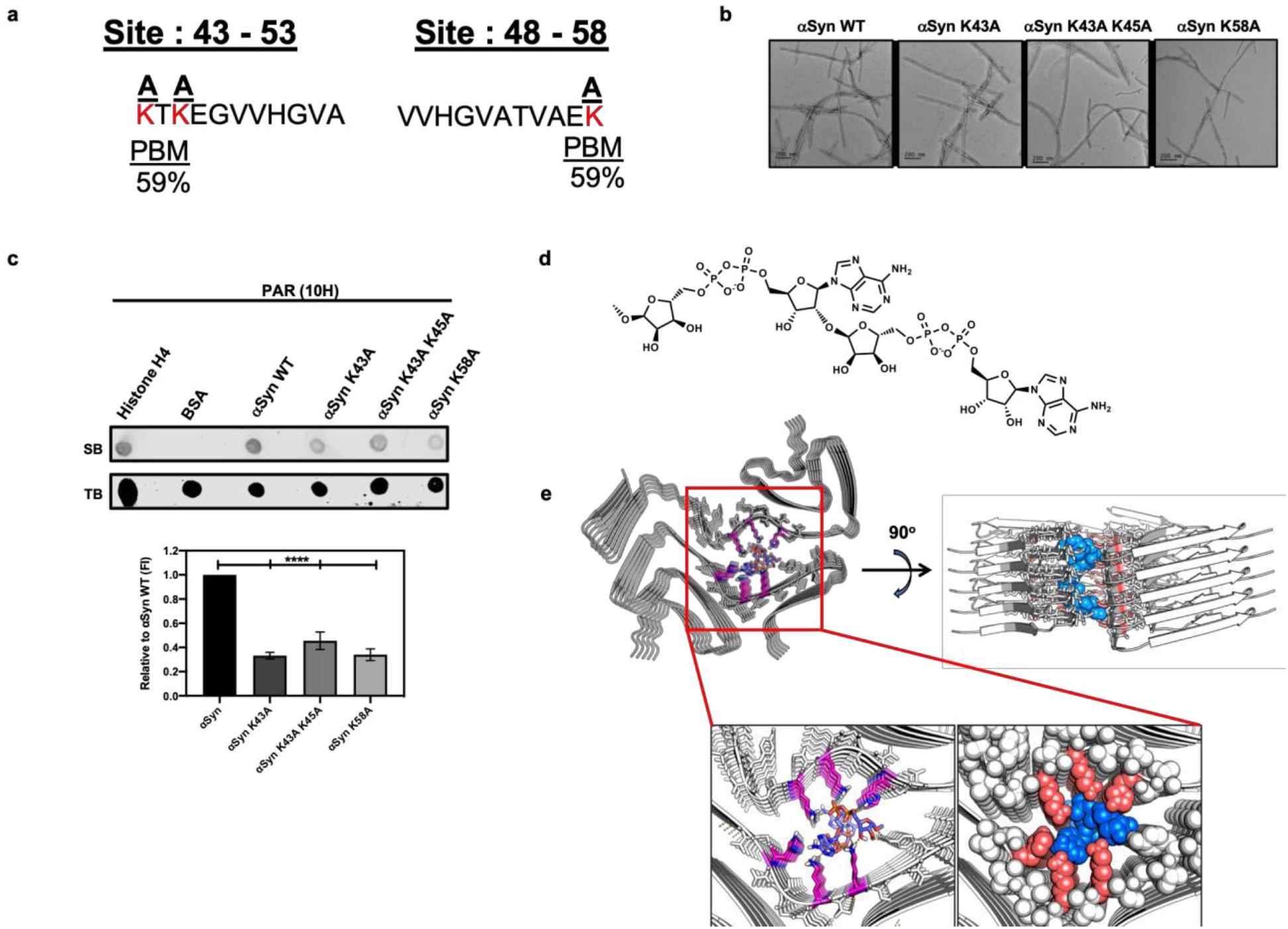
PAR interacts with αSyn via electrostatic forces in the N-terminal region of the protein. (a) Alignment of the full αSyn sequence with PBMs yielded a 59% PAR-binding probability at amino acid residues 43-54 and 48-58 on αSyn. Lysine amino acid residues (red) at the two sites were substituted to neutral alanine residues. (b) Three mutants of αSyn were generated with compromised PAR-binding sites. Two of the αSyn mutants had a point mutation at amino acid residues K43 & K58 respectively, while the third mutant had two point mutations at positions K43 and K45. All mutants were fully fibrillated within 72 hr. Scale bar 200 nm. (c) PAR immunodot blot (top), whereby WT and mutant αSyn fibrils were spotted onto a membrane, along with, histone H4 (positive control), and BSA (negative control) and incubated with PAR polymer to assess PAR binding. Semi-quantitative analysis (bottom) of WT and mutant αSyn fibril signal intensity normalized to WT αSyn signal. One-way ANOVA (n =3). ****P < 0.0001. (d) Chemical structure of PAR-dimer used in molecular docking studies. (e) Cryo-EM structure of MSA Type I αSyn fibril interacting with the PAR-dimer complex with a low free binding energy of −15.6 kcal/mol.

To better characterize αSyn-PAR interactions, we substituted positively charged lysine residues at the two PAR-binding sites predicted by the NPS@PATTINPROT tool (Fig. 6a). Using site-directed mutagenesis, we generated three mutants of αSyn with compromised PAR-binding sites by replacing lysine residues with neutral alanine residues (Fig. 6a). Two of the αSyn mutants had a single point mutation at amino acid residues K43 & K58 respectively, while the third mutant had two point mutations at positions K43 and K45 (Fig. 6a,b). Fibrillization of the αSyn mutants was followed using Thioflavin T (ThT) (Extended Data Fig. 4a) and transmission electron microscopy (TEM) (Fig. 6b). The introduced mutations decreased the rate of αSyn fibril formation by increasing the lag-phase relative to the wild type (WT) protein (Extended Data Fig. 4a). Despite the initial slow aggregation kinetics, all mutants were fully fibrillated within 72 h as shown by ThT (Extended Data Fig. 4a) and TEM (Fig. 6b). We observed kinetic variability in mutant-seeded formation of αSyn fibrils from the single point mutant K43 and the double point mutant K43 and K45, as determined by ThT (Extended Data Fig. 4b). Cytotoxicity assays on IMR-5 neuroblastoma cells revealed that the αSyn mutant fibrils were less toxic compared to PAR-bound αSyn WT fibrils (Extended Data Fig. 4c), these results are in accordance with previous observations^28^ which show that PAR-bound fibrils display higher neurotoxicity. Thus, reinforcing the notion that PAR enhances the toxicity of αSyn aggregates.

To test if PAR binding was affected in the generated αSyn mutant fibrils, we performed PAR-binding immunodot blot (Fig. 6c). WT and mutant αSyn fibrils were spotted onto a membrane along with a PAR-binding protein, histone H4 (positive control) and BSA (negative control). Incubation with PAR polymer, followed by immunoblotting with a PAR-specific antibody (10H), revealed that both histone H4 and αSyn WT fibril bound to PAR. Interestingly, we observed a decrease in PAR binding for all three mutants when compared to αSyn WT fibril (Fig. 6c).

To assess if the decrease in PAR binding on the alanine mutant fibrils was a direct result of substituting positively-charged lysine residues –and not just due to the introduction of a point mutation, we performed an additional PAR-binding immunodot blot with αSyn A53T fibril. From this, we observed that PAR binds the αSyn A53T mutant with similar signal intensity compared to αSyn WT fibril (Extended Data Fig. 3d). Based on these results, we confirm that PAR binding to αSyn is primarily mediated by electrostatic interactions at positions 43-58 of the N-terminal region. Our data also indicates that PAR binds a known familial point mutation of αSyn (A53T) (Extended Data Fig. 4d). The latter has direct relevance in patients who harbor the A53T variant of αSyn – as this variant has been shown to be aggregation prone and has been directly linked to autosomal dominant early onset PD.

### PAR, MSA, and beyond

The cryo-EM structure of Sarkosyl-insoluble αSyn filaments isolated from five MSA cases was recently reported^61^. Two different filament types were reported, type I and type II. Both filament types consist of two different protofilaments having an extended N-terminus and compact C-terminal body. In addition, the interface between the two different protofilaments consist of a non-proteinaceous density in the region of K43 and K45 in one protofilament, and K58 of the other protofilament. Since replacement of these lysine residues with an alanine residue diminished PAR binding (Fig. 6c) and given that PAR and pαSyn interact in both PD/PDD (Fig. 5a,c) and MSA brain tissue (Extended Data Fig. 3c), we propose that this non-proteinaceous density is likely to be PAR.

To evaluate this hypothesis, a series of computational chemistry studies (Fig. 6d,e) were conducted in order to probe the interaction of a PAR-dimer (Fig. 6d) with the cryo-EM structure of αSyn, specifically the type I filament^61^. From this, we found a very strong fit of the PAR-dimer in the space occupied by this non-proteinaceous density – with strong ionic interactions between these lysine residues and the diphosphate moiety of the PAR-dimer and hydrogen bond interactions between tyrosine-39 and histidine-50 with the adenine group and ribose groups (Fig. 6e). Altogether, our results are consistent with PAR being the non-proteinaceous density in the αSyn filaments reported in MSA.

## Discussion

PAR is a highly branched polymer that has been best characterized as a recruiter of DNA repair factors during single-strand DNA break repair. However, in recent years the role of PAR outside of the nucleus has become clearer and the role of this polymer in neurodegeneration stands as a promising avenue for better understanding the molecular basis of neurotoxicity that gives way to neurodegeneration. Specifically, looking at PAR in PD points to a potential role for this polymer as a nucleating driver of αSyn aggregation. Understanding this interaction and relating it to disease progression points to PAR and presumably PAR-αSyn interactions, specifically, as potential early biomarkers for disease. Such a finding, holds wide implications for early prognosis and further presses on the need to develop robust PARP inhibitors that reduce PARP-1 activity without eliciting toxicity in cells. In addition, the significant interactions between PAR-pαSyn observed in post mortem PD/PDD patient samples suggests that this interaction might be prevalent in disease. Our studies also confirm that PAR and αSyn interact via electrostatic forces involving positively-charged lysine residues on αSyn. Altogether, our observations potentiate the need to further study the association of αSyn-PAR as it pertains to disease and explore the possibility of using this interaction as the basis for developing disease modifying therapies and prognostic tools to track disease progression. Our findings reinforce the notion that PARP-1 hyperactivity (and resulting intracellular PAR accumulation) is a key driver of neurodegeneration. Thus, highlighting the need to evaluate if early interventions with PARP-1 inhibitors can translate into preventative measures for individuals who are at higher risk of developing PD.

## Study Design

The objectives of this study were to evaluate the role of PAR in driving monomeric αSyn aggregation, elucidate PAR binding to pαSyn in pathology-associated cell, murine, and human post mortem brain samples, and identify the amino acid residues involved in αSyn fibril-PAR binding. These controlled laboratory experiments involved the use of immunostaining, PLA, molecular biology, and computational chemistry techniques.

Immunostaining and PLA results were analyzed using CellProfiler 3.0^62^ software, whereby, specialized pipelines were implemented to identify and count PAR and pαSyn staining (Cell/particle counting pipeline) or PLA dot signal (modified Speckle Counting pipeline; whereby PLA dots were identified within cells’ cytoplasm). These measurements were then normalized to DAPI count to obtain an output measure of *total target signal/total cell* count per field of view (10 – 15) for each sample.

For animal studies, sample size for each age group was n = 3. Age groups were determined via PAR immunostaining, whereby, mice that were 12 mo of age and older displayed higher PAR intensity. The oldest mice in our study were between 17 mo and 18 mo of age, therefore, we defined 17 mo as the end point for our murine data collection. Mice that were older than 12 mo were included in the “aged” group, whereas, mice that were younger than 12 mo were included in the “young” group. Similarly, we used littermate controls for the “aged” group to account for age-related effects. Information on the strain, sex, and age of the mice used in this study can be found in Supplementary Table 2. For studies on post mortem tissue samples, PD/PDD cases were characterized by PD type pathology (Supplementary Table 3).

## Materials and Methods

### PAR polymer

Purified PAR polymer chains (commercially obtained from TREVIGEN) were synthesized from PARP-1 in the presence of NAD^+^, cleaved and subsequently purified. PAR chain lengths ranged in size from 2-300 ADP-ribose subunits, with a final concentration of 10 μM.

### Cell Culture

SH-SY5Y-αSyn cells were transfected using Lipofectamine 2000 (Invitrogen) with pcDNA3.1 expression vector following manufacturer’s protocol. The expression vector contained the full-length human wild type αSyn cDNAs, cloned in the polylinker region at the KpnI and ApaI sites. Stable transfected cell lines were selected and maintained in complete medium containing 300 μg/ml G418 (Invitrogen) ^63^. The cells were maintained in DMEM/F12 media with GlutaMAX supplement (Thermofisher scientific, Cat#10565018), 10% heat-inactivated fetal calf serum (FBS), 100 units/ml penicillin and 100 μg/ml streptomycin (Pen-Strep), in a humid atmosphere of 5% CO^2^ and 95% O^2^ at 37 °C.

### *BioPORTER* Experiments

SH-SY5Y-αSyn cells were seeded at concentrations of 16,000 cells/well in Nunc® Lab-Tek® Chamber Slide™ system (8 wells, 0.8 cm^2^/well) (Millipore®, C7182-1PAK) for fluorescent microscopy experiments (IF and PLA) or at 2 × 10^6^ in 150 mm TC treated (ThermoFisher Scientific, Cat#FB012925) dishes for biochemical assays (Western, ELISA, immunodot) 24 h before incubation with either PAR + *BioPORTER,* ADP-HDP *+ BioPORTER* or *BioPORTER* alone (vehicle control). *BioPORTER* Protein Delivery Reagent “QuikEase Kit” (Genlantis, Cat#BP502424) was prepared according to the manufacturer’s protocol, briefly described as follows. Either PAR or ADP-HDP were diluted in 100 μL PBS to a final concentration of 50 nM, the diluted solution was then added to a QuikEase Single-Use Tube containing the dried BioPORTER reagent, mixed by pipetting 3-5 times, incubated at room temperature (RT) for 5 min and gently vortexed (post-incubation) for 3-5 s. Opti-MEM I Reduced Serum Medium (Life Technologies Inc., Cat#31985062) was used to bring the final volume in each QuikEase Single-Use Tube to 500 μL. The cells were washed once with Reduced Serum Medium 1 h before *BioPORTER* delivery, then replenished with either 200 μL (chamber slide cells) or 6 mL (150 mm dish cells) of Opti-MEM I (ThermoFisher Scientific, Cat#31985062). *BioPORTER* medium mix was added at a 1:1 volume ratio in the cells grown in chamber slides and a 1:3 volume ratio in the cells grown in 150 mm dishes. The cells were subsequently incubated for 4 h at 37 °C. After 4 h, one volume of 20% serum-containing medium was added directly to the chamber slides/dishes, 24 h post-*BioPORTER* delivery, the medium was aspirated from the chamber slides/dishes and replenished with complete growth medium (DMEM/F12 media with GlutaMAX supplement). 48 h after *BioPORTER* delivery, the cells were washed 2X with PBS and processed for downstream experiments.

### Cell Viability Assays

Cells were seeded in black wall clear bottom 96-well plates (Corning®) at concentrations of 1000 cells/well. For PAR toxicity assay, SH-SY5Y cells were treated with three concentrations of PAR ranging from 25 to 100 nM for 48 h. For pre-formed fibril (PFF) toxicity assays, IMR-5 cells were treated with 750 nM PFF for five days. Following treatment incubation, the cells were assayed for viability using the luminescent based assay, CellTiter Glo (Promega Corp.), following manufacturer’s protocol. Plates were read on an Enspire multimode plate reader (PerkinElmer, Inc.). Data was normalized to percent survival at each concentration evaluated by dividing the luminescent signal in treated wells by the average of PBS controls. Experiments were repeated three times.

### Transfection of Hela cells with EGFP-αSyn-A53T

Hela cells were transiently transfected using Lipofectamine 3000 (Invitrogen) with a pEGFP-C1 mammalian vector containing full length human A53T variant αSyn with a fusion EGFP tag (Addgene, Plasmid #40823), following the manufacturer’s protocol. The transfected cells were maintained in complete medium (DMEM media, 10% FBS, 1% Pen-Strep) and used within 2 days of transfection for downstream experiments.

### MitoSOX™ Red

Hela-EGFP-αSyn-A53T cells were seeded at concentrations of 5,000 cells/well in Nunc® Lab-Tek® Chamber Slide™ system. *BioPORTER* delivery was used to deliver 50 nM PAR polymer, as described above, Bio*PORTER* alone was used as a vehicle control. 48 h post-PAR delivery, cells were treated with 75nM MitoSOX™ Red reagent (ThermoFisher, Cat#M36008) for 30 min at 37 °C to detect production of superoxide by mitochondria in Hela-EGFP-αSyn-A53T cells. After incubation, the cells were fixed with ice cold methanol for 15 min at 4°C, washed 3X with PBS and processed for imaging using Zeiss Axio Widefield microscope (20x/0.8).

### Animals

M83-SNCA*A53T mice expressing human A53T variant αSyn were obtained from The Jackson Laboratory, Bar Harbor, ME (JAX stock #004479). All mice were on B6;C3H genetic background. Animals were housed under controlled temperature and lighting conditions and had free access to food and water. All animal procedures were approved by IACUC and were in accordance with the National Institutes of Health Guide for the Care and Use of Laboratory Animals.

### Human Post Mortem Brain αSyn Pathology Analysis

Human brain samples were obtained from University of Pennsylvania’s Center for Neurodegenerative Disease Research Brain Bank and were evaluated with standardized histopathological methods as described^64–66^.

### Immunofluorescence (IF) Staining

48 h post-*BioPORTER* delivery, SH-SY5Y-αSyn cells (seeded on chamber slides at 16,000 cells per well) were fixed on ice with 4% paraformaldehyde for 8 min. The cells were then washed 3X with PBS and permeabilized with 0.1% Triton X-100 for 10 min at RT. After permeabilization, the cells were washed 3X with PBS-T (PBS with 0.1% Tween-20) at RT. After the third wash, 200 μL of 10% goat serum (ThermoFisher, Cat#50062Z) was added to each well for 1 h at 37°C to block non-specific immuno binding. After blocking, the cells were sequentially incubated with primary antibodies (Supplementary Table 1) targeting PAR (10H) and pαSyn (ps129) overnight at 4°C. Following primary antibody incubation, the cells were washed 3X with PBS-T. After the third wash, the cells were then sequentially incubated with secondary antibodies (Supplementary Table 1) for 1 h at 37°C, washed 3X with PBS-T and stained with DAPI. Coverslips were placed on each slide and the slides were allowed to dry overnight at 4°C. Images were captured using Zeiss LSM 710 confocal (40x/1.4 Oil) and Zeiss Axio Widefield (20x/0.8) microscopes (Supplementary Table 4).

Human post mortem tissue sections were treated with TrueBlack (TrueBlack® Lipofuscin Autofluorescence Quencher) according to manufacturer’s protocol, in order to eliminate lipofuscin autofluorescence before immunostaining.

Following the blocking step with 10% goat serum, murine tissue sections underwent an additional blocking step with anti-mouse IgG (Supplementary Table 1) for 1 h at 37°C in order to reduce nonspecific signal from secondary antibodies directed against PAR antibody (10H), which is a mouse monoclonal primary antibody.

### Proximity Ligation Assay (PLA)

48 h post-*BioPORTER* delivery, chamber slides cells (SH-SY5Y-αSyn cells) were processed with regards to fixation and permeabilization using the IF protocol described in the previous section. In situ PLA was performed according to the manufacturer’s protocol, briefly described as follows. Following permeabilization, cells were blocked using Duolink® Blocking Solution for 1 h at 37°C. PAR primary antibody (Supplementary Table 1) was diluted in Duolink® Antibody Diluent, added to the cells and incubated overnight at 4°C. Following overnight incubation with PAR primary antibody, the cells were washed 2X with Duolink® Wash Buffer A, then incubated with Duolink® PLA Probe (goat anti-mouse *MINUS*) for 1 h at 37°C. After incubation with PLA Probe *MINUS,* the cells were washed 2X with Wash Buffer A, blocked with Duolink® Blocking Solution for 1 h at 37°C and incubated overnight at 4°C with primary antibody targeting pαSyn (Supplementary Table 1). Following overnight incubation, the cells were washed 2X with Wash Buffer A and incubated with Duolink® PLA Probe (goat anti-rabbit *PLUS*) for 1 h at 37°C. Following the sequential addition of primary antibodies and corresponding PLA Probes, the cells were processed with respect to ligation (Duolink® Ligation buffer and Ligase), amplification (Duolink® Amplification buffer and Polymerase) and imaging using Zeiss Axio Widefield (20x/0.8) microscope. Human post mortem tissue sections were treated with TrueBlack (TrueBlack® Lipofuscin Autofluorescence Quencher) according to manufacturer’s protocol, in order to eliminate lipofuscin autofluorescence before PLA. Murine tissue sections underwent an additional blocking step with anti-mouse IgG (Supplementary Table 1) for 1 h at 37°C to reduce nonspecific signal from goat anti-mouse *MINUS*.

### Western Blotting

Cells were washed twice with cold PBS and scraped into ice cold 1X lysis buffer (Thermo Scientific, RIPA buffer) containing 1X protease and phosphatase inhibitors (Thermo Scientific, Halt™ Protease and Phosphatase Inhibitor Cocktail) and allowed to lyse, while rocking, at 4°C for 1 h. Lysates were then cleared by centrifugation at 13,000 rpm for 20 min. Supernatant was collected, and the protein was quantified using BioRad DC protein quantification assay. All samples were then diluted to a final concentration of 2 μg/μL with 1X Laemmli buffer. Samples were separated on 4-20% BioRad TGX pre-packed gels at 100 V for 1 h. Gels were transferred to a PVDF membrane using BioRad turbo transfer at 1.3 A for 7 min. Next, membranes were washed 4X in PBS with 0.2% tween 20 and incubated in Odyssey Blocking buffer (Li-COR), 0.2% tween-20, and 0.1% SDS for 1 h. Membranes were incubated overnight at 4 °C with Primary antibodies (Supplementary Table 1) and detected with fluorescent secondary antibodies (IRDye, Supplementary Table 1). Uniform regions of interest were applied to each lane to calculate total fluorescence intensity, which was representative of total target protein. Either Histone H3 (loading control) or Revert 700 stain (Revert 700 Total Protein, Li-COR) were used to calculate final relative protein expression for each lysate. Following Revert 700 stain, membranes were washed 2X with Revert 700 wash buffer (Li-COR) for 5 min each. Membranes were imaged using Li-COR ODYSSEY CLx scanner. Experimental data represent the average of 3 independent experiments.

### ELISA

A Human αSyn oligomer ELISA kit (MyBioSource, MBS730762) was used to quantify of αSyn oligomer levels from cell samples treated with either PAR + *BioPORTER* or *BioPORTER* alone for 48 h. Briefly, cells were washed 2X with PBS, scraped into ice cold 1X PBS, centrifuged at 1000 rpm for 5 min and resuspended in 100 μL of PBS. The resuspended cells were then lysed using a hand held sonicator for three pulses at 40% amplitude. 1X protease and phosphatase inhibitors were added to the lysate following sonication; the samples were kept on ice and immediately processed for protein quantification using DC protein quantification assay. All samples were diluted to a final concentration of 2.5 μg/μL and loaded into coated wells from the ELISA kit all steps were followed according to the manufacturer’s protocol.

### A11 Immunodot Blot

2 μL of cell lysates (2.5mg/mL) were placed on a nitrocellulose membrane and allowed to air dry at RT for 1 h. At this point, the membrane was incubated in Revert 700 protein stain for 10 min at RT and washed 2X with Revert 700 wash buffer for 5 min each. The membrane was imaged using Li-COR ODYSSEY CLx scanner at 700 nm to visualize total protein. After imaging, the membrane was washed for 5 min in Revert 700 reverse solution (Li-COR) in order to remove protein stain. The membrane was blocked using Odyssey blocking buffer for 1 h at RT and then incubated overnight at 4°C with A11, oligomer-specific antibody (Supplementary Table 1). Following overnight incubation, the membrane was washed 3X using PBS-T (0.1% Tween-20). Secondary Antibody (Supplementary Table 1) was then added to the membrane and incubated at RT for 1 h. After incubation, the membrane was washed 3X in PBST and processed for imaging.

### αSyn Protein Expression and Purification

Protein expression and purification was done following previously published protocol^67^. Briefly, the plasmid encoding the human αSyn sequence was transformed into Escherichia coli BL21(DE3) and the cells were grown on agar/LB plates with ampicillin (100 μg/mL) overnight at 37°C. The next day a single colony was inoculated into 100 mL Luria-Bertani (LB) containing ampicillin (100 μg/mL). The culture was incubated at 37°C overnight with shaking at ~200 rpm. The following day, 10 mL of the overnight culture was diluted with 1 L of LB media supplemented with ampicillin and this culture was incubated at 37°C until OD600 reached 0.6 – 0.7. Protein expression was induced by addition of isopropyl-β-D-thiogalactoside (IPTG) to a final concentration of 1 mM and continued to grow at 18°C overnight. After induction, cells were harvested by centrifugation at 4°C (20 min, 4,000 g). The typical yield of wet-cell paste was 2 g/L. Cells were suspended in a lysis buffer (5 mL for 1 g of cell paste) containing 25 mM Tris, 20 mM imidazole, 50 mM NaCl (pH 8) with a protease inhibitor (phenylmethylsulfonylfluoride, 0.5 mM final concentration and protease inhibitor cocktail from Cell Signaling Technology). Cells were lysed by sonication on ice for 10 min (20 s on, 20 s off). The crude cell lysate was then centrifuged at 20,000g for 30 min, and the supernatant was mixed with Ni-NTA resin (Clontech, 3 mL) and kept on a rocker at RT for 30 min. The resin was then washed with 100 mL wash buffer (25 mM Tris, 20 mM imidazole, 50 mM NaCl, pH 8). The protein was eluted with a buffer containing 25 mM Tris, 300 mM imidazole, 50 mM NaCl (pH 8). Fractions containing the protein were identified by UV-Vis spectroscopy, combined and was treated with β-mercaptoethanol (200 mM final concentration) overnight at RT to cleave the C-terminal intein. The next day, the protein was concentrated to 3 mL and dialyzed against buffer containing 25 mM Tris, 50 mM NaCl, pH 8. After dialysis, the protein mixture was loaded onto Ni-NTA column and the pure αSyn protein was collected in the flow through fractions. The combined protein fractions were concentrated and dialyzed against buffer containing 50 mM Tris, 150 mM NaCl, pH 7.5. The purity of the protein was confirmed by SDS-PAGE. Protein concentration was determined by measuring the absorbance at 280 nm and using the calculated (ExPASy) extinction coefficient of 5960 M-1cm-1.

### Site-directed Mutagenesis

αSyn mutations were generated by performing site directed mutagenesis using the following primers:

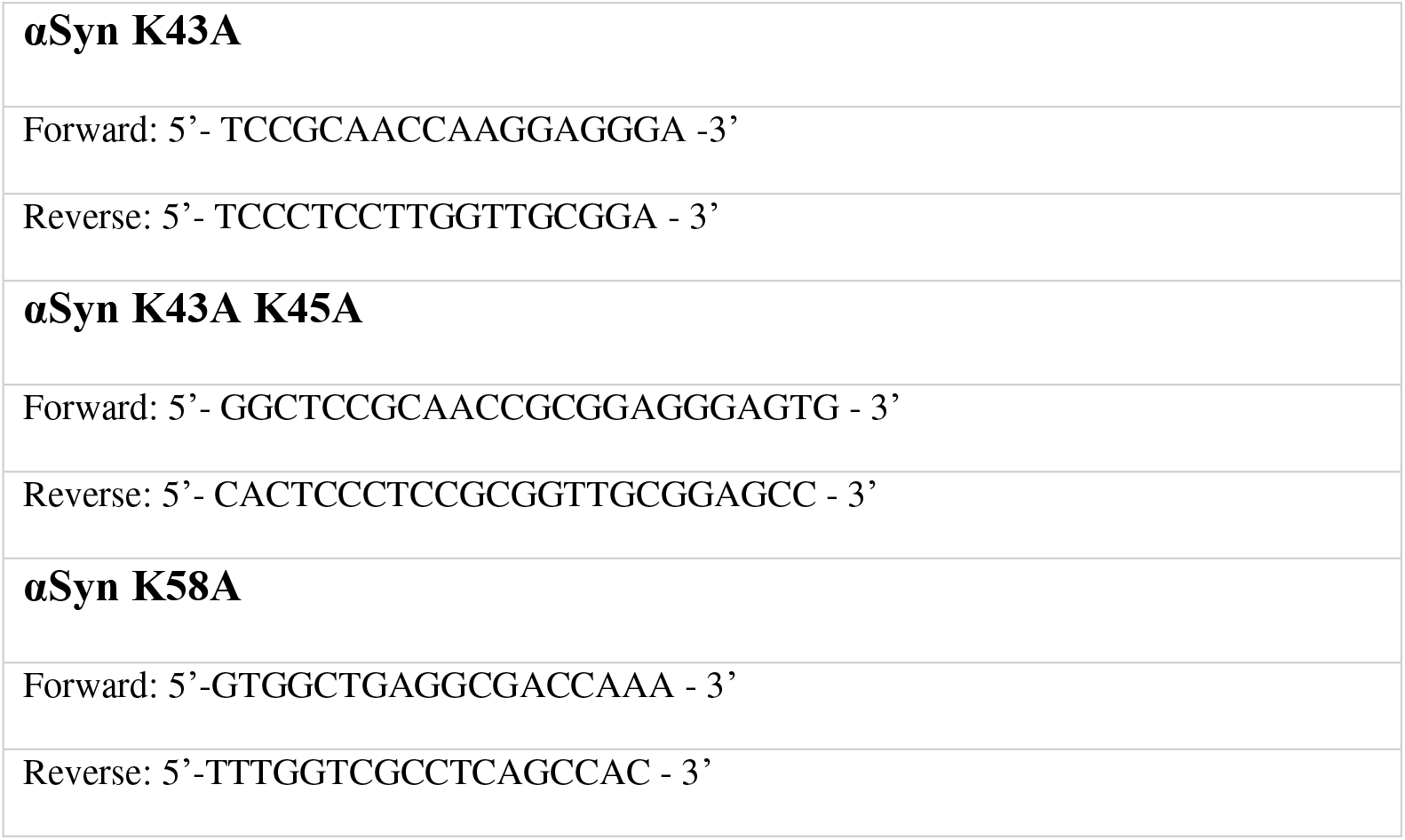

All plasmids and inserts were sequenced and confirmed to be free of any errors.

### PAR Binding Motifs (PBM)

*hxbxhhbbhhb* (h are hydrophobic residues, b are basic residues, and x is any amino acid residue)^22^. [HKR]-X-X-[AIQVY]-[KR]-[KR]-[AILV]-[FILPV]^22,60^.

### Preparation of αSyn fibrils

Purified αSyn monomer (100 μM) was incubated in buffer containing 50 mM Tris (pH 7.5), 150 mM NaCl and 0.05% NaN3 for 72 h at 37 °C with shaking at 1000 rpm in a Fisher Scientific Mixer. Fibrils with poly(ADP-ribose) (PAR) were prepared in the presence of 100 nM PAR polymer (TREVIGEN) following the procedure described above.

### Transmission electron microscopy (TEM)

The 100 μM fibril stock solution was diluted 4X with water and samples (5 μL) were spotted onto glow-discharged formvar/carbon-coated, 200-mesh copper grids (Ted Pella). After 1 min, grids were washed briefly with water and stained with one 10 μL drop of 2% w/v uranyl acetate for 1 min. The excess stain was removed by filter paper and the grids were dried under air. Samples were imaged with a Tecnai FEI T12 electron microscope at an acceleration voltage of 120 kV. Images were recorded on a Gatan OneView 4K Cmos camera.

### PAR Immunodot blot

PAR-binding motif (PBM) were identified by aligning the PBM consensus to αSyn using the PATTINPROT search engine (NPS@PATTINPROT). For immunodot analysis, either 1 μg fibrils, Histone H4 (positive control), or bovine serum albumin (negative control) were blotted onto a 0.2 μm nitrocellulose membrane (Biorad). Membranes were left to dry for 60 min, then incubated in DPBS supplemented with 0.05% Tween-20 (PBS-T) for 10 min. The membrane was then incubated with 50 nM PAR polymer in PBS-T for 2 hours with rocking at RT. The membrane was washed 5 X with PBS-T (5 min each) and blocked with PBSMT (5% milk in PBS-T) for 2 hours at RT. After the blocking step, the membrane was incubated in primary antibody (Supplementary Table 1) in PBS-T at 4 °C overnight. After 5 washes in PBSMT (5 min each), the membrane was incubated with secondary antibody (Supplementary Table 1) in PBSMT for 1 hour at RT. The membrane was washed 3X in PBSMT, 2 X in PBS-T and 2X in DPBS (5 min each). The membrane was then imaged using Li-COR ODYSSEY CLx scanner. Spot intensities were measured using Image Studio software. Revert 700 protein stain was used for total protein staining measurement. Blotted membranes were incubated with protein stain for 5 min, rinsed with Revert 700 wash buffer, and imaged using Li-COR ODYSSEY CLx scanner.

### Thioflavin-T (ThT) assay

Fluorescence spectra were obtained on an EnSpire Multimode Plate Reader (Perkin Elmer), using 440 nm as excitation wavelength. Samples were prepared by mixing protein and ThT in buffer (50 mM Tris, 150 mM NaCl, pH 7.4) in a 96-well plate to a final concentration of 100 μM and 10 μM, respectively. The plate was shaken at 1000 rpm at 37 °C and fluorescence emission was measured at 490 nm over time.

### Molecular docking

The PAR-dimer structure used in our studies was based on Lambrecht et al.^68^ and drawn on ChemDraw Profession 15.1 (PerkinElmer Informatics, Inc.). It was then imported to Chem3D Ultra 15.1 (PerkinElmer Informatics, Inc.) to minimize the PAR-dimer by MMFF94 force field for preparation of molecular docking. Molecular docking studies were performed via AutoDock 4.269 plugin on PyMOL (pymol.org). Cryo-EM structure of MSA Type I αSyn fibril (PDB ID 6XYO, Resolution 2.6 Å) was obtained from RCSB Protein Data Bank (www.rcsb.org). Polar hydrogens were added to the fibril structure. Non-polar hydrogens were removed from the PAR-dimer. A grid box with a dimension of 30 × 30 × 30 Å^3^ was applied to the MSA Type I αSyn fibril structure covering the non-proteinaceous density pocket at the protofilament interface^70^.

The Lamarckian Genetic Algorithm with a maximum of 2,500,000 energy evaluations was used to calculate 100 αSyn fibril-PAR binding poses. The αSyn fibril-PAR complex with the most contacts and low free binding energy was reported.

### Quantification and Statistical Analysis

All measurements were taken from distinct samples. Data points in each graph are mean (± SEM); where “n” indicates the number of biological replicates for each experiment. T-tests, one-way ANOVA, and two-way ANOVA followed by Tukey’s post hoc test were performed and are described in each figure legend. Statistical significance was set at P < 0.05. All statistical analyses were carried out using Graphpad prism 8 software.

## Acknowledgments

SH-SY5Y-αSyn cells were a gift from Dr. Harry Ischiropoulos, University of Pennsylvania. The plasmid encoding the human αSyn sequence was a gift from Dr. James Petersson, University of Pennsylvania.

## Author contributions

L.N.P. performed all the cell-based studies and ex vivo animal and patient experiments, along with the computational protein alignment. Z.L.Z. helped produce all the purified proteins and fibrils used in this project. Z.L.Z performed TEM, ThT, and PAR immunodot blot analysis. L.N.P. and Z.L.Z. designed the primers for mutagenesis. J.Y.L. maintained cell cultures and aided in the experimental set-up for PLA. C.J.H. performed molecular docking studies. M.E.S. maintained αSyn protein expression and purification. K.J.E. provided assistance with PLA and transfection of Hela cells with EGFP-αSyn-A53T. K.C.L. provided support with animal model and experimental design. V.M.Y.L. and J.Q.T. provided support with experimental design and characterization of human post mortem brain tissue from PD/PDD and non-PD patients.

## Data and materials availability

All data generated and analyzed in this study are included in this published article and its extended data files.

---

**Extended Data Fig 1.**
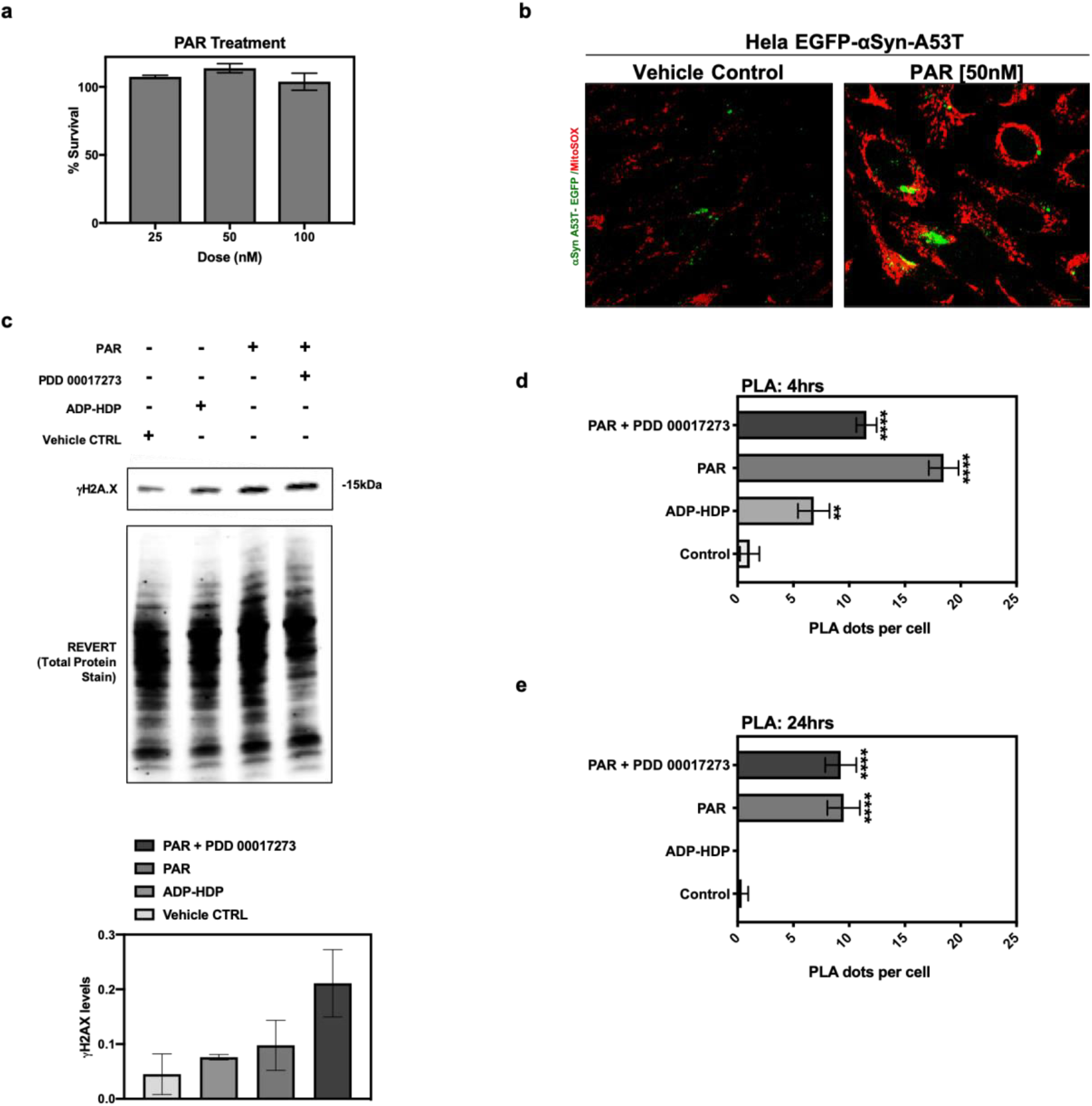
Intracellular effects of PAR on αSyn. (a) Cytotoxicity assay on SH-SY5Y neuroblastoma cells treated in triplicate with three different concentrations of PAR polymer for 48 h. (b) Hela cells transfected with EGFP-αSyn-A53T and treated with PAR (50 nM) for 48 h; representative ROIs from merge channel images showing αSyn A53T-EGFP (green) and increased superoxide levels i.e. higher red intensity, in the PAR treated vs. *BioPORTER* alone (vehicle control) cells. Images were captured using Zeiss Axio Widefield (20x/0.8) microscope. Experiments were performed in triplicate. (c) Representative Western blot and graph of γH2AX levels in PAR, PAR + 1 μM PDD00017273 (PARGi) and ADP-HDP treated SH-SY5Y-αSyn cells. Experiments were repeated independently three times (n =3). (d and e) Quantification data from a time course PLA on SH-SY5Y-αSyn cells treated with PAR, PAR + 1 μM PDD00017273 (PARGi) or ADP-HDP for 4 h (d) and 24 h (e). Bars represent means ± SEM. Two-way ANOVA followed by Tukey’s post hoc test (n =3). ****P < 0.0001.

**Extended Data Fig 2.**
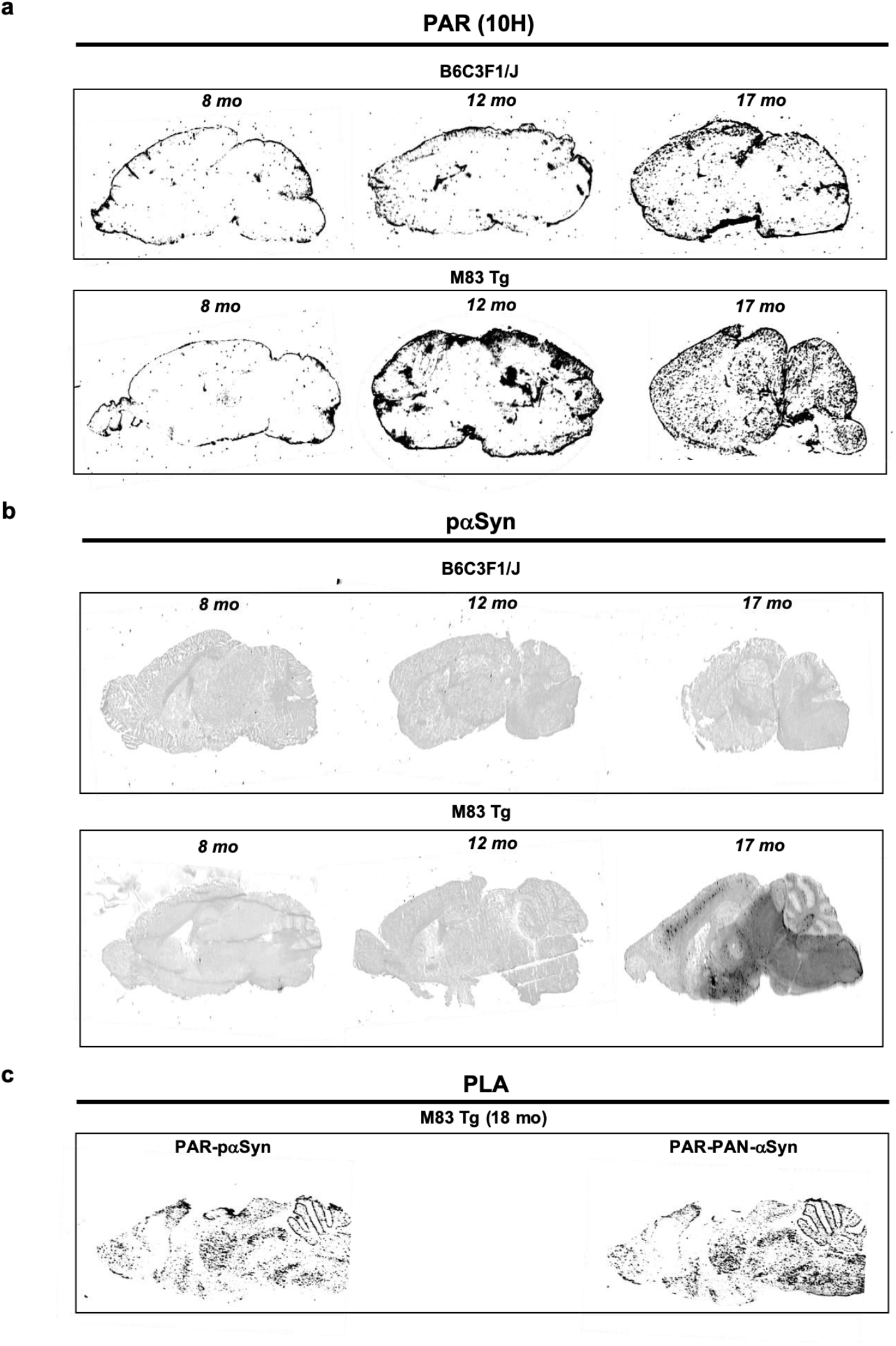
Characterization of PAR and pαSyn expression in murine brain sections. Representative images of endogenous PAR (a) and pαSyn (b) levels in sagittal brain sections from B6C3F1/J (top panel) and M83 Tg (bottom panel) mice at three different age groups (8 mo, 12 mo, and 17 mo). (c) PLA on PAR-pαSyn (left) and PAR-PAN-αSyn (right) sagittal brain sections from M83 Tg mice at 18 mo of age. All sections were 10 μm thick and all images were captured using a Li-COR ODYSSEY CLx scanner.

**Extended Data Fig 3.**
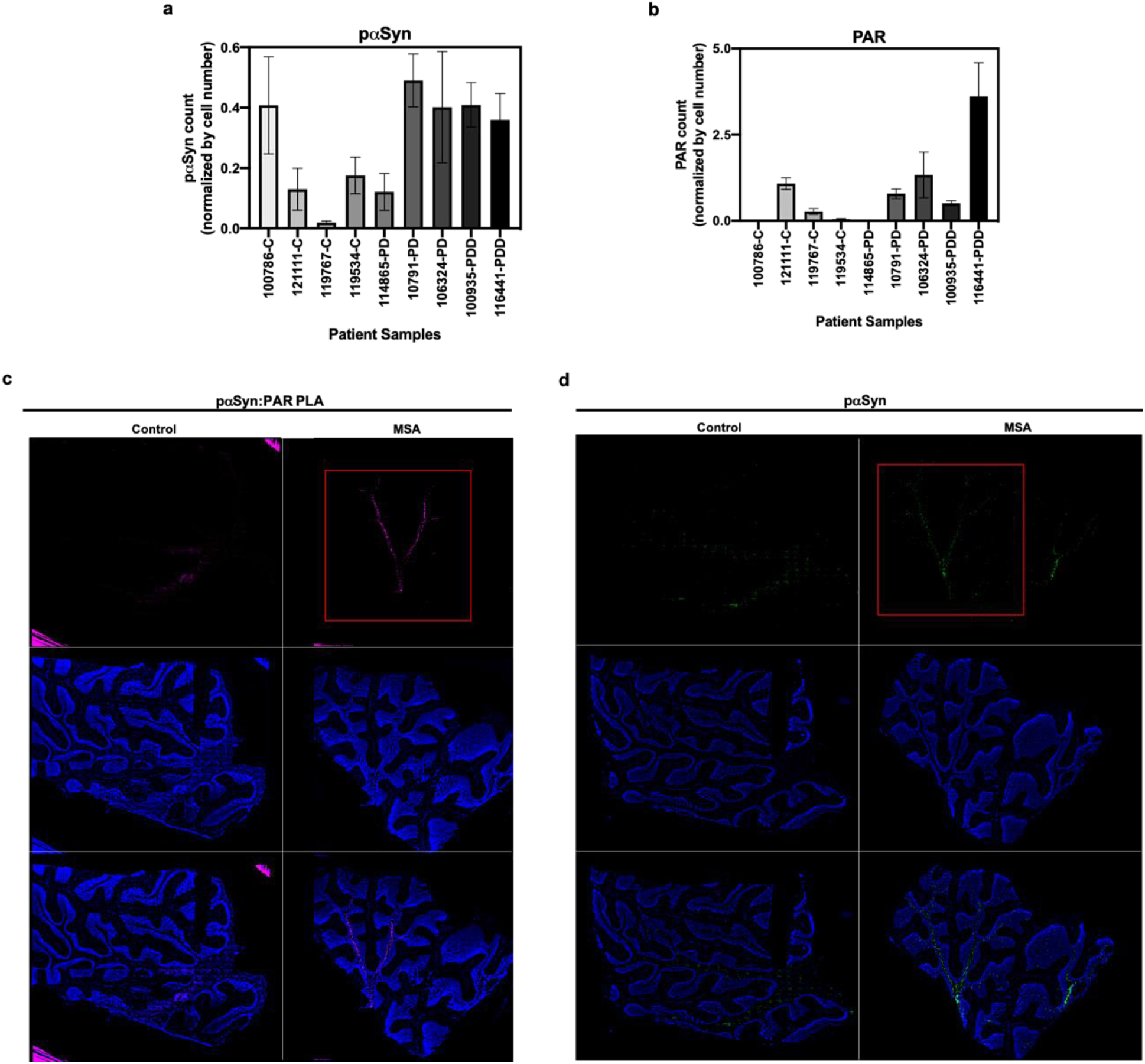
Characterization of PAR and pαSyn expression in post mortem brain sections. (a and b) Immunostain quantification of pαSyn and PAR expression for all human PD/PDD and non-PD post mortem brain samples used in this study. (c) PLA on cerebellum sections from control and MSA patients showing PLA (top panel), DAPI (middle panel), and merge (bottom panel) channel images. (d) Standard immunostain of adjacent cerebellum sections from control and MSA patients showing pαSyn (top panel), DAPI (middle panel), and merged (bottom panel) channel images. Red boxes indicate PLA signal (c) and matching pαSyn IF (d) in adjacent MSA tissue sections. Images were captured using Zeiss Axio Widefield (20x/0.8) microscope.

**Extended Data Fig 4.**
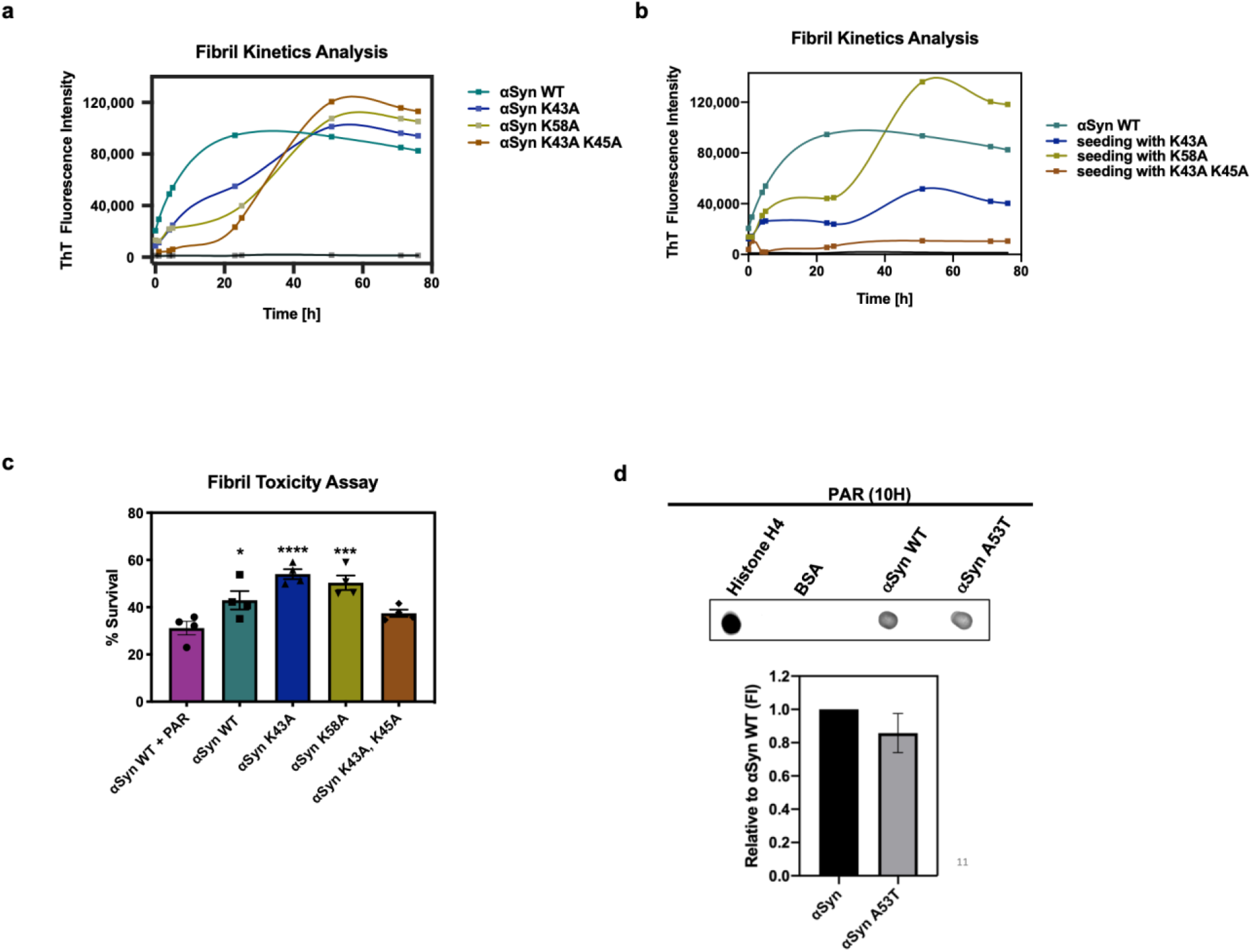
Characterization of αSyn fibril and PAR binding. (a) ThT kinetic analysis and characterization of the three point mutants generated via site-directed mutagenesis against αSyn WT protein. (b) Fibril seeding variability of the three point mutants as determined by ThT kinetic analysis. (c) PFF cytotoxicity data on neuroblastoma cell line IMR-5. Cells were treated for 5 days with 750 nM PFF. Bars represent means ± SEM. Two-way ANOVA followed by Tukey’s post hoc test (n = 4). *P < 0.03, ***P < 0.0005, ****P < 0.0001. (d) Representative PAR immunodot blot to assess PAR-αSyn A53T binding. Signal intensity was normalized to αSyn WT signal. Histone H4 and BSA were used as positive and negative controls for PAR binding respectively.

## Supplementary Information

**Supplementary Table 1.** Antibody information.

| Antibody | Antigen | Host Species | Dilution | Source |
| --- | --- | --- | --- | --- |
| A11 | oligomers | Rabbit | 1:10,000 (DotBlot) | ThermoFisher |
| Syn303 | αSyn (target N-terminus, conformation specific) | Mouse | 1:500 (WB) | Biologend |
| γH2AX | γH2AX | Mouse | 1:1000 (WB) | MilliporeSigma |
| Histone 3 (H3) | Histone 3 | Rabbit | 1:1000 (WB) | CST |
| 81A | aSyn (Phosphorylation at Ser-129) | Mouse | 1:1,000 (IF; cells) | Abcam |
| αSyn (PS129) | αSyn (Phosphorylation at Ser-129) | Rabbit | 1:500 (IF, PLA) | Abcam |
| PAR (10H) | PAR | Mouse | 1:500 (IF, PLA; cells), 1:1000 (IF, PLA; tissue), 1:5,000 (DotBlot) | Enzo |
| Goat F(ab) anti-Mouse IgG (H&I) | Mouse IgG | Mouse | 1:100 (IF & PLA; murine tissue) | Abcam |
| αSyn (PAN-αSyn) | αSyn (targets C-terminus, polyclonal) | Rabbit | 1:500 (PLA) | Invitrogen |
| IRDye 800 Goat anti-Mouse | Mouse IgG | Mouse | 1:5,000 (PAR DotBlot) | LI-COR Biosciences |
| Goat anti-Mouse IgG Alexa Fluor 800 | Mouse IgG | Mouse | 1:10,000 (WB) | Invitrogen |
| Goat anti-Rabbit IgG Alexa Fluor 680 | Rabbit IgG | Rabbit | 1:10,000 (WB, A11 DotBlot) | Invitrogen |
| Goat anti-Rabbit IgG Alexa Fluor 568 | Rabbit IgG | Rabbit | 1:400 (IF) | Invitrogen |
| Goat anti-Mouse IgG Alexa Fluor 489 | Mouse IgG | Mouse | 1:400 (IF) | Invitrogen |
| PLA Probe Anti-Rabbit PLUS | Rabbit IgG | Rabbit | 1:5 (PLA) | MilliporeSigma |
| PLA Probe Anti-Mouse MINUS | Mouse IgG | Mouse | 1:5 (PLA) | MilliporeSigma |

**Supplementary Table 2.** Animal Information.

| **Strain** | **Sex** | **Age** | **Group** |
| --- | --- | --- | --- |
| M83 SCNA*A53T | M | 6 | M83 Tg young |
| M83 SCNA*A53T | M | 7 | M83 Tg young |
| M83 SCNA*A53T | M | 8 | M83 Tg young |
| M83 SCNA*A53T | M | 12 | M83 Tg aged |
| M83 SCNA*A53T | M | 15 | M83 Tg aged |
| M83 SCNA*A53T | M | 17 | M83 Tg aged |
| <sup>1</sup>B6C3F1/J | M | 12 | B6C3F1/J aged |
| <sup>1</sup>B6C3F1/J | M | 12 | B6C3F1/J aged |
| <sup>1</sup>B6C3F1/J | M | 17 | B6C3F1/J aged |
| <sup>1</sup>non-Tg mice (littermate control) |  |  |  |

**Supplementary Table 3.** Patient Information.

| **INDDID** | **Clinical Group** | **Age at Death** | **Sex** | **Region** |
| --- | --- | --- | --- | --- |
| 100786 | Control | 61 | M | Striatum |
| 119767 | Control | 83 | M | Striatum |
| 119534 | Control | 72 | F | Striatum |
| 121111 | Control | 68 | M | Middle Frontal Gyrus |
| 114865 | PD | 62 | M | Striatum |
| 107091 | PD | 72 | F | Striatum |
| 100935 | PDD | 80 | M | Striatum |
| 106324 | PD | N/A | N/A | Middle Frontal Gyrus |
| 116441 | PDD | 82 | M | Middle Frontal Gyrus |

**Supplementary Table 4.**
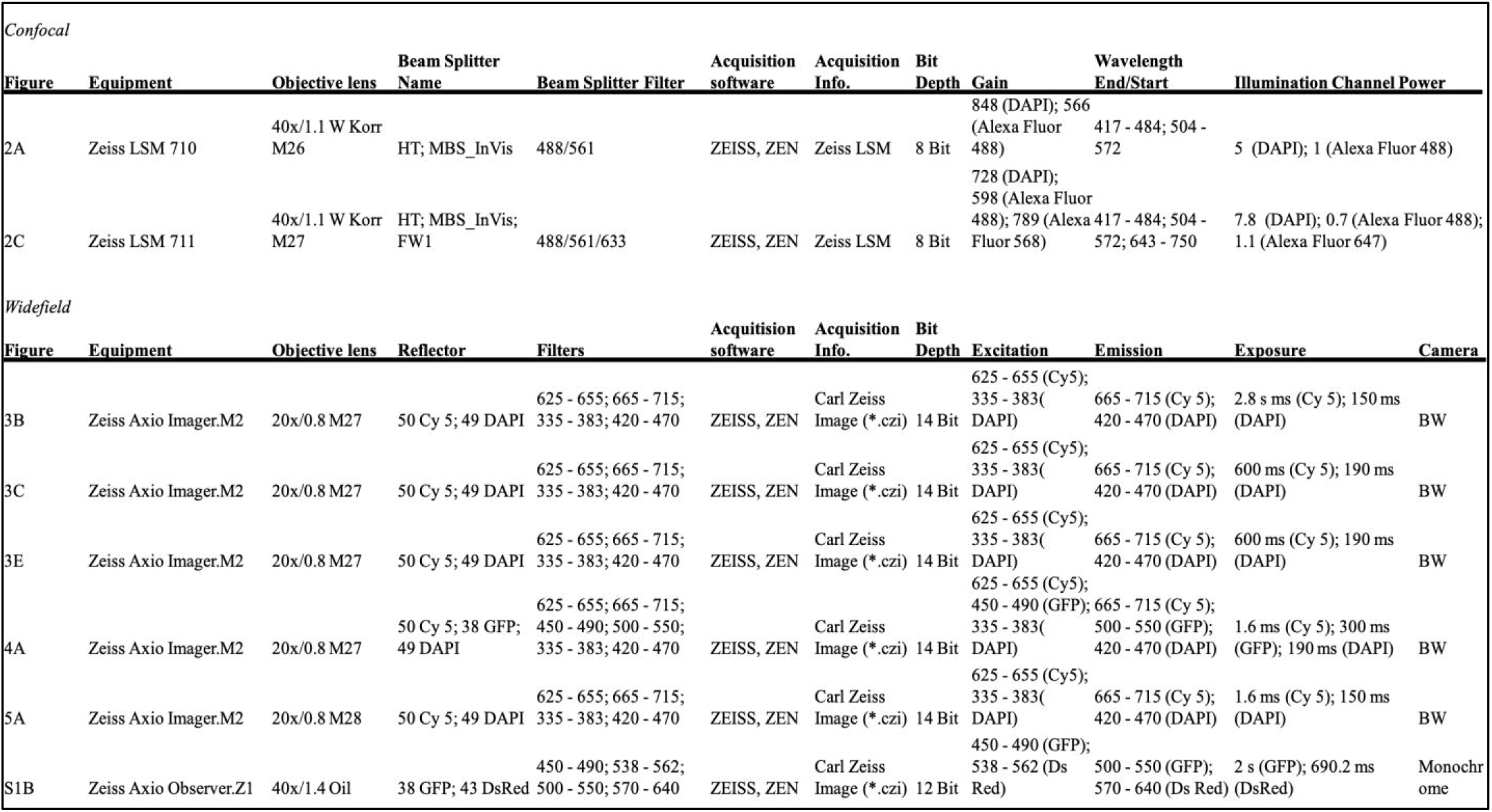
Microscopy Settings.

